# The Crk adapter protein is essential for *Drosophila* embryogenesis, where it regulates multiple actin-dependent morphogenic events

**DOI:** 10.1101/654558

**Authors:** Andrew J. Spracklen, Emma M. Thornton-Kolbe, Alison N. Bonner, Alexandru Florea, Peter J. Compton, Rodrigo Fernandez-Gonzalez, Mark Peifer

## Abstract

Small SH2/SH3 adapter proteins regulate cell fate and behavior by mediating interactions between cell surface receptors and downstream signaling effectors in many signal transduction pathways. The Crk family has tissue-specific roles in phagocytosis, cell migration and neuronal development, and mediates oncogenic signaling in pathways like that of Abelson kinase. However, redundancy among the two mammalian family members and the position of the *Drosophila* gene on the fourth chromosome precluded assessment of Crk’s full role in embryogenesis. We circumvented these limitations with shRNA and CRISPR technology to assess Crk’s function in *Drosophila* morphogenesis. We found Crk is essential beginning in the first few hours of development, where it ensures accurate mitosis by regulating orchestrated dynamics of the actin cytoskeleton to keep mitotic spindles in syncytial embryos from colliding. In this role, it positively regulates levels of the Arp2/3 complex, its regulator SCAR, and F-actin in actin caps and pseudocleavage furrows. Crk loss leads to loss of nuclei and formation of multinucleate cells. We also found roles for Crk in embryonic wound healing and in axon patterning in the nervous system, where it localizes to the axons and midline glia. Thus, Crk regulates diverse events in embryogenesis that require orchestrated cytoskeletal dynamics.

## Introduction

Cells have an incredible ability to receive a wide-array of environmental inputs and respond in context-dependent manners during embryonic development and adult tissue homeostasis. Loss of these responses results in developmental defects and contributes to cancer. Cells receive many different signals, including growth factors, adhesion to neighbors, extracellular matrix engagement, or bioactive lipids. These inputs lead to diverse cellular responses, including changes in transcription, proliferation, cell shape, motility, or cell death. Despite the diversity of inputs and the responses they evoke, different pathways often share an underlying logic. Signals are usually received by plasma membrane receptors (e.g., receptor tyrosine kinases (RTKs), integrins, or cadherins), and information is transduced throughout the cytoplasm by a cadre of downstream effectors, often initiated by formation of multiprotein complexes associated with the cytoplasmic tails of transmembrane receptors.

Small adapter proteins, such as Grb2, Crk, or Nck, play critical roles in both signal amplification and specificity (e.g. (Lettau *et al*., 2009; Belov and Mohammadi, 2012; Hossain *et al*., 2012; Martinez-Quiles *et al*., 2014)). These adapters confer signal specificity by using Src homology domain 2 (SH2) and 3 (SH3) or other protein interaction domains to form physical links between activated cell surface receptors and their downstream effectors. A particular small adapter protein only interacts with select receptors and effectors, ensuring a specific response. Small adapters can also amplify cell signaling.

CT10 Regulator of Kinase (Crk) proteins are a conserved family of small modular adapters shared by diverse animal lineages (reviewed in Feller, 2001; Birge *et al*., 2009; Hossain *et al*., 2012). In vertebrates, there are two genes encoding three protein isoforms: CrkI, CrkII (alternative splice forms of *crk*) and Crk-like (CrkL). An N-terminal SH2 domain allows them to bind tyrosine phosphorylated proteins, while a canonical SH3 domain binds proline-rich motifs (“PXXP”). CrkII and CrkL have an additional, non-canonical C-terminal SH3 domain that does not bind PXXP motifs, but serves important regulatory roles. This organization allows Crk family adapters to link PXXP-containing downstream effectors to upstream proteins containing phosphorylated tyrosines, like activated RTKs.

Our interest in Crk was initiated by our work on the developmental roles of the non-receptor tyrosine kinase Abelson (Abl; reviewed in Khatri *et al*., 2016). Abl is the oncogene activated in chronic myelogenous leukemia, and the target of the clinically effective kinase inhibitor imatinib (Greuber *et al*., 2013). *Drosophila* Abl plays important roles in many processes, ranging from the cytoskeletal rearrangements of syncytial development and cellularization (Grevengoed *et al*., 2003), to cell shape changes during gastrulation and dorsal closure (Grevengoed *et al*., 2001; Fox and Peifer, 2007), to roles in axon outgrowth and guidance in the central nervous system (e.g. Gertler *et al*., 1989; Bashaw *et al*., 2000; Kannan and Giniger, 2017). When we examined Abl’s mechanisms of action, we found that while kinase activity is important, an even more essential determinant of Abl function is a short, conserved PXXP motif in the Abl linker known in mammals to bind Crk, Nck, and Abi (Rogers *et al*., 2016). Crk is a known Abl substrate and a potential upstream activator (Hossain *et al*., 2012), and Crk knockdown alleviates the effects of Bcr-Abl transformation (e.g. Seo *et al*., 2010). This stimulated our interest in Crk as a potential Abl regulator or effector in *Drosophila*.

Crk family adapters modulate a wide-range of cell behaviors in cultured cell models, including cell growth, migration, adhesion, and immune function (reviewed in Feller, 2001; Birge *et al*., 2009; Hossain *et al*., 2012). For example, in fibroblasts, CrkI/II and CrkL play redundant roles in regulating cell growth and proliferation (Park *et al*., 2016), cell morphology and motility (Park and Curran, 2014; Park *et al*., 2016), and oncogene-mediated cell transformation (Koptyra *et al*., 2016). Crk proteins localize to focal adhesions to promote their stability (Senechal *et al*., 1996; Li *et al*., 2003; Antoku and Mayer, 2009; Watanabe *et al*., 2009; Park and Curran, 2014), as well as promote cell spreading and motility (Chodniewicz and Klemke, 2004). Crk proteins also work downstream of integrins to coordinate motility and substrate mechanosensing in T-cells (Roy *et al*., 2018). Not surprisingly, Crk family proteins also appear to play important roles in cancer, promoting cell transformation and tumor growth (reviewed in Sriram and Birge, 2010; Kumar *et al*., 2014).

While work in cultured cells provided numerous insights, Crk’s roles in normal development are less well understood. CrkII is dispensable for normal development (Imaizumi *et al*., 1999). Mice lacking both CrkI and CrkII are inviable and display multiple defects in cardiovascular and craniofacial development (Park *et al*., 2006). CrkL knockout mice exhibit reduced viability and have multiple developmental defects associated with cardiac and neural crest derivatives (Guris *et al*., 2001), at least in part through effects on FGF signaling (Moon *et al*., 2006). Interestingly, CrkL maps to a chromosomal region commonly deleted in DiGeorge syndrome patients, and phenotypic similarities suggest its loss may contribute to this disease. Tissue specific knockouts revealed additional roles for Crk or CrkL in places ranging from T cell migration (Huang *et al*., 2015; Roy *et al*., 2018) to Bcr-Abl transformation of mouse hematopoietic progenitors (Seo *et al*., 2010). The two mammalian Crk family members have redundant roles in some tissues. Simultaneous neural-specific knockout of both Crk and CrkL revealed essential overlapping roles downstream of Disabled-1 in the Reelin pathway regulating neuronal positioning (Park and Curran, 2008). Similar overlapping roles were seen in the kidney (George *et al*., 2014) and lens (Collins *et al*., 2018). However, to our knowledge, there has been no analysis of mouse embryos lacking both CrkI/II and CrkL from the onset of development.

In *C. elegans* and zebrafish Crk adapters play important developmental roles in cell migration, axon outgrowth, phagocytosis, and myoblast fusion. The *C. elegans* Crk homolog Ced-2 is required for phagocytic clearance of apoptotic cells, cell migration, and motor neuron development, as part of a Crk/DOCK180/Rac pathway (Reddien and Horvitz, 2000; Gumienny *et al*., 2001; Wu *et al*., 2001; Wu *et al*., 2002). In zebrafish, morpholino-mediated knockdown of either Crk or CrkL, both of which are broadly expressed, impairs fusion of fast-twitch myoblasts (Moore *et al*., 2007), while overexpressing Crk or CrkL enhances fusion.

*Drosophila* has a single Crk family member, thus providing a simplified model to define core functions of Crk. In the fly, biochemical studies revealed Crk can directly bind potential upstream regulators like the receptor tyrosine kinase PVR (Ishimaru *et al*., 2004) and the Ig-family receptor Sticks and Stones (Kim *et al*., 2007), as well as potential downstream effectors like the DOCK-class Rac Guanine-nucleotide exchange factor Myoblast City (Mbc; *Drosophila* DOCK180; (Galletta *et al*., 1999; Balagopalan *et al*., 2006)), Verprolin 1 (*Drosophila* WASP-interacting protein (Kim *et al*., 2007)), and its upstream regulator Blown fuse (Jin *et al*., 2011).

Much of this analysis has focused on roles of Crk in mediating Dock180/ELMO/Rac1 signaling to the actin cytoskeleton via WASP, SCAR, and the Arp2/3 complex (reviewed in Deng *et al*., 2017). Tissue-specific RNAi revealed a role for Crk upstream of Mbc/Elmo/Rac1 signaling in phagocytic clearance of axonal debris (Ziegenfuss *et al*., 2012) and of pathogenic protein aggregates by glial cells (Pearce *et al*., 2015). Crk also acts in the Mbc/Elmo/Rac1 engulfment pathway in developmental pruning of larval neurites (Tasdemir-Yilmaz and Freeman, 2014), removal of apoptotic neurons (Nakano *et al*., 2019), and salivary gland clearance by autophagy (McPhee *et al*., 2010). *Drosophila* Crk also plays important roles in phagocytic uptake of pathogenic bacteria (Elwell *et al*., 2008; Pielage *et al*., 2008), similar to roles Crk plays in mammalian cells (reviewed in Martinez-Quiles *et al*., 2014).

The Dock180/ELMO/Rac1 pathway also acts downstream of the RTK PVR during thorax closure and border cell migration, and tissue-specific RNAi and use of a hypomorphic *crk* allele support roles for Crk in these processes (Ishimaru *et al*., 2004; Geisbrecht *et al*., 2008). Another DOCK family GEF, the DOCK3 relative Sponge, acts in parallel to Mbc in both processes (Bianco *et al*., 2007; Morishita *et al*., 2014). Dock180/ELMO/Rac1 signaling also regulates the actin cytoskeleton during myoblast fusion, but despite clear evidence that Crk directly binds several proteins essential for myoblast fusion, it is unclear whether Crk is required for this event. Overexpressing a membrane-targeted form of Crk results in myoblast fusion defects (Abmayr *et al*., 2003), but Mbc mutants lacking all Crk binding sites fully restore myoblast fusion (Balagopalan *et al*., 2006). Crk may play a role in muscle development, however, as a muscle-specific RNAi led to a flightless phenotype, with missing indirect flight muscles, (Schnorrer *et al*., 2010). Thus, tissue specific Crk knockdown revealed several important developmental roles, many involving DOCK/ELMO/Rac regulation of actin.

While informative, these analyses via tissue specific RNAi do not reveal the core, conserved roles for Crk family adapters during embryogenesis. We sought to generate embryos devoid of Crk, and thus analyze its full set of functions. Despite its identification nearly 20 years ago, there has been surprisingly little genetic analysis of *Drosophila crk*, largely due to the fact that *crk* maps to the fourth chromosome, which has been less genetically tractable. The only existing allele was a hypomorphic mobile element insertion (Ishimaru *et al*., 2004). Modern tools allowed us to circumvent this issue. We used second-generation shRNA lines and developed a suite of genetic tools, including a null allele, a powerful gene replacement platform, and overexpression constructs. These tools allowed us to manipulate Crk levels in ways not previously possible, revealing Crk is required for embryonic viability and plays critical roles during actin-dependent morphogenic events.

## Results

### Crk is required for embryonic viability

Despite Crk’s importance in oncogenic signaling and the known tissue specific roles of the two mammalian family members, its full importance in normal development remains unclear. *Drosophila* provides a superb model to examine this role, as there is only a single family member, thus avoiding the partial redundancy of the mammalian proteins. However, the position of *crk* on the fourth chromosome hindered the ability to examine its function, with previous studies limited to the use of a probable hypomorphic allele derived from P-element insertion (Ishimaru *et al*., 2004) or tissue-specific RNAi. New tools offered the opportunity to develop ways to reduce or eliminate Crk function.

We took two approaches: using CRISPR to generate tagged and null alleles, and RNAi to knockdown function. We generated a *crk* null allele, crk^ΔattP^, using CRISPR to replace the entire *crk* locus with a 50 base-pair attP phage recombination site and positive selectable marker (Fig 1A). This allele was verified by PCR (Suppl. Figure 1A,B). Consistent with the previously characterized allele *crk^KG00336^*(Ishimaru *et al*., 2004), homozygous zygotic *crk*^Δ*attP*^ mutants were embryonic viable but died as pupae, revealing *crk* to be an essential gene. We next created a series of rescue constructs, in which either the complete genomic region or an mNeonGreen-tagged cDNA were targeted back into the locus at the attP site (Fig 1B). The genomic rescue construct and one version of the cDNA rescue constructed were flanked by FRT sites (Fig 1B; full descriptions are in Suppl. Fig. 1C), allowing deletion via the FLP recombinase, to facilitate production of females whose germlines were homozygous mutant. We also generated a Crk antibody, using the full-length protein as the antigen, which recognize a single protein of molecular weight ∼31 kDa (Fig 1C), consistent with the predicted size of Crk (31kD). All three rescue constructs are expressed at essentially normal levels (Fig. 1C), and all rescued viability and fertility.

**Figure 1.**
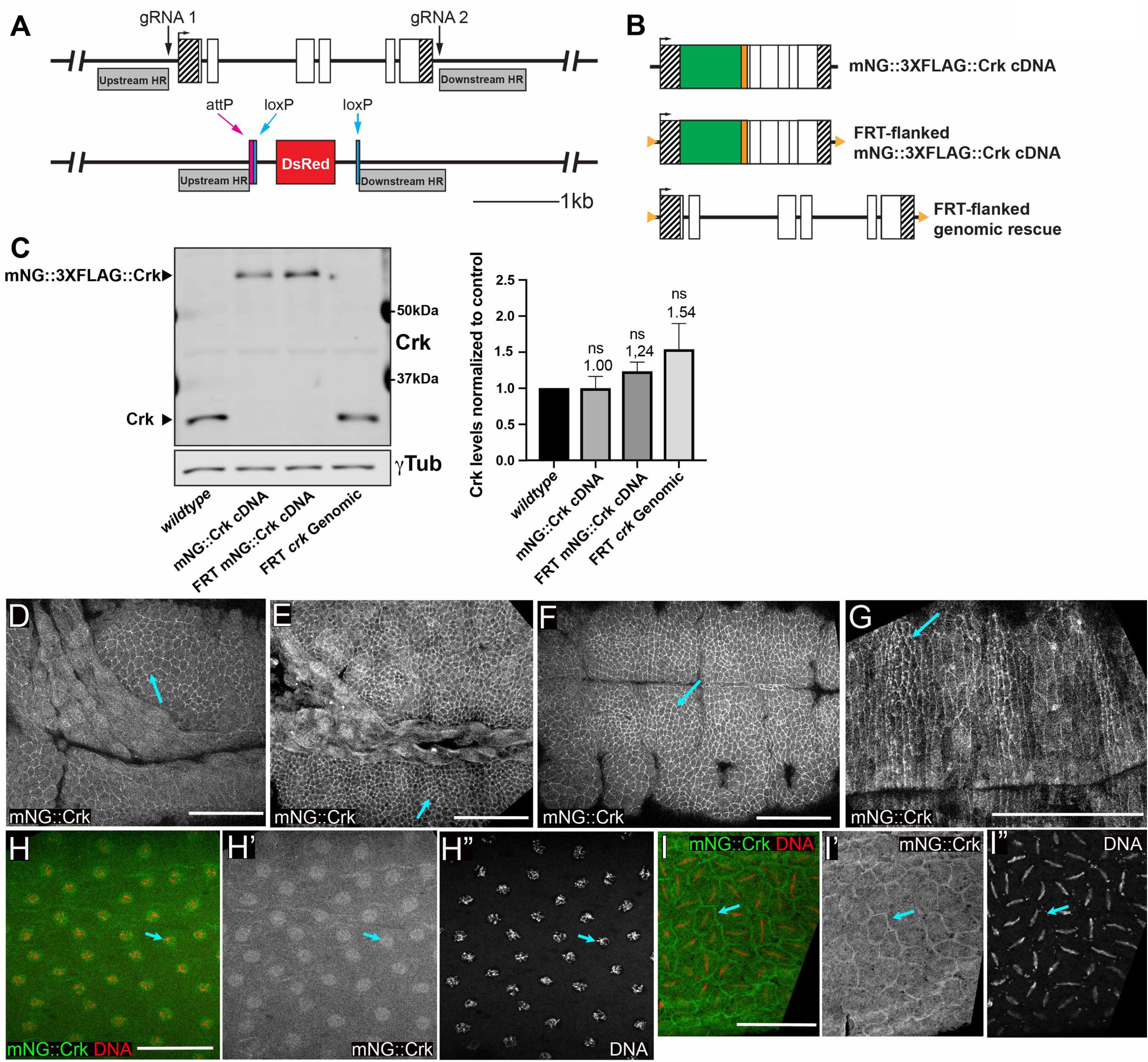
Generation of a *crk* null allele and a gene replacement platform. A. Schematic representation, *crk* locus before and after CRISPR/Cas9-mediated deletion of *crk* to generate *crk^ΔattP^*. We used two guide RNAs (gRNAs) to introduce double-strand breaks 110bp upstream of the transcription start site and 99bp downstream of the end of the 3’UTR. Using a double-stranded donor repair template containing flanking homology regions (indicated by shaded grey bars), we replaced the *crk* coding sequence with an attP recombination site and a loxP-flanked, positive visual marker (3xP3-DsRed expressed in the eye). B. Schematic representation, rescue alleles targeted into *crk^ΔattP^*. The three constructs include: cDNA-based constructs containing a tandem N-terminal tag (monomeric NeonGreen (mNG; green) and a 3XFLAG tag (orange)), with or without flanking FRT sites (orange triangles), and an FRT-flanked genomic rescue construct carrying the exact sequence removed in generating *crk^ΔattP^*. C. Representative Western blot and quantification of Crk levels in wildtype and *crk^ΔattP^*-targeted rescue lines. Rescue constructs restore endogenous levels of Crk; n=4, error bars=SEM, ns=not statistically significant. D-I. Representative images, Crk localization throughout embryonic development. Embryos expressing mNeonGreen-tagged Crk (mNG::Crk) inserted at endogenous locus. The tagged protein was imaged using mNG fluorescence. H and I were also stained to reveal DNA. D. Stage 9. E. Stage 10. F. Stage 11. G. Stage 14. After gastrulation Crk localized to a cytoplasmic pool with enrichment at the cell cortex (arrows). H,I. Syncytial blastoderm. Crk is enriched in actin caps that overlie nuclei (H, arrow) and at pseudocleavage furrows (I, arrow). Scale bars= 50µm.

While zygotic mutants were embryonic viable, we suspected this was due to maternally contributed Crk, as *crk* mRNA is maternally loaded and broadly expressed (Galletta *et al*., 1999). We hypothesized Crk would be broadly expressed and essential for embryonic morphogenesis; consistent with this, our mNeonGreen-tagged construct targeted into the endogenous locus was expressed throughout embryogenesis (Fig. 1D-I). After gastrulation we observed cortical enrichment in the ectoderm and developing epidermis, along with a cytoplasmic pool (Fig 1D-G, arrows)—this continued until at least the completion of dorsal closure (Fig 1G).

To directly test the hypothesis that Crk is essential for embryonic development, we used RNAi to maternally and zygotically deplete *crk*. We used two independent UAS-driven shRNA lines targeting different regions of *crk*, developed by the Transgenic RNAi Project (TRiP; Perkins *et al*., 2015). shRNA constructs can mimic maternal/zygotic loss of gene products, when expressed using germline drivers (Staller *et al*., 2013). We drove expression of shRNAs targeting *crk* using two copies of a strong germline driver (*maternal αtubulin*-GAL4; *matα*-GAL4) to knockdown maternal *crk*. Both RNAi lines impair embryonic viability, but to differing extents. The stronger of the two lines, referred to below as *crkS-RNAi*, severely impaired embryonic viability, reducing viability to 1.2% (Fig. 2A; n=922), while the second line, referred to below as *crkW-RNAi*, resulted in weaker effects, reducing viability to 71.4% (Fig. 2A; n=960), compared to UAS-shRNA only controls, which exhibited 97.9% (Fig. 2A; n=711) and 97.5% (Fig. 2A; n=770) viability, respectively.

**Figure 2.**
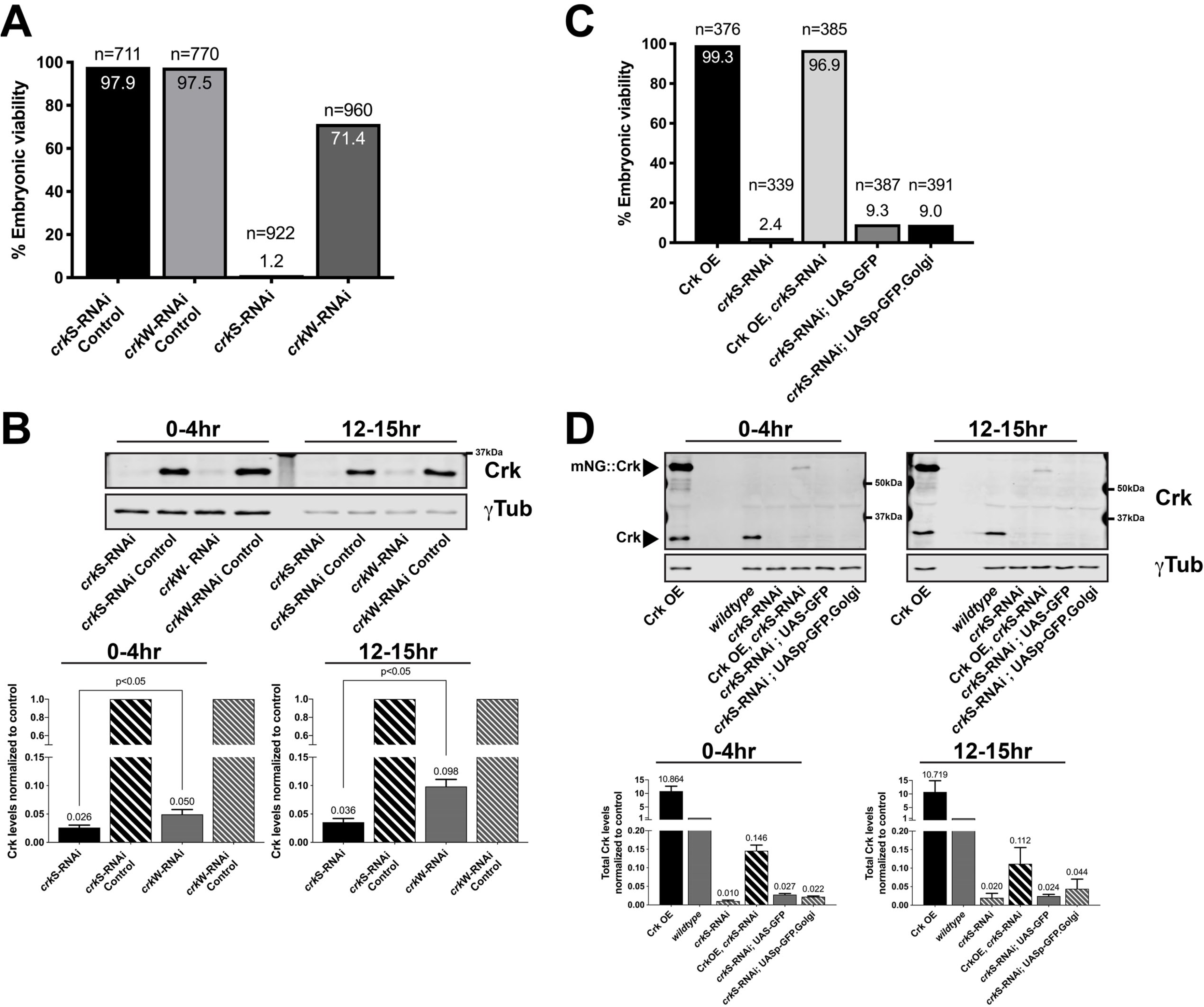
Crk is required for embryonic viability. A. Quantification of embryonic viability following shRNA-mediated knockdown of Crk. Compared to shRNA-only controls (*crkS-RNAi* Control and *crkW-RNAi* Control), Crk knockdown using either of two shRNA lines driven by a pair of strong maternal GAL4 drivers reduces embryonic viability. B. Representative Western blot and quantification, Crk levels in Crk knockdown embryos compared to controls. *crkS-RNAi* embryos have a stronger reduction in Crk levels compared to *crkW-RNAi* embryos; n=4, error bars=SEM. C. Quantification of embryonic viability following overexpression of Crk or simultaneous knockdown/overexpression of Crk. Simultaneous overexpression of Crk fully restores embryonic viability in *crkS-RNAi* embryos. Crk overexpression (Crk OE; ∼10 fold higher than endogenous) alone has no impact on embryonic viability. Expressing two different extraneous GFP-tagged proteins to control for effects of two UAS promotors does not substantially rescue lethality. D. Representative Western blot and quantification of Crk levels in embryos overexpressing Crk (lane one), wildtype (lane four, note lanes two and three intentionally left empty), *crkS-RNAi* emrbyos (lane five), undergoing simultaneous knockdown/overexpression (lane six), or in *crkS-RNAi* embryos simultaneously expressing GFP-tagged proteins to control for effects of two UAS promotors (lanes seven and eight), n=3, error bars-SEM.

To verify the shRNA constructs knocked down *crk*, and, in parallel, to test specificity of our antibody, we examined Crk accumulation at two stages of embryogenesis by immunoblotting. Both shRNAs strongly reduced Crk protein levels during early embryonic development, with *crkS-RNAi* dropping levels to 2.6% and *crkW-RNAi* to 5.0% at 0-4hr (Fig. 2B; left), consistent with their differential effects on embryonic lethality. Expression remained strongly reduced at 12-15 hrs (3.6% and 9.8%, respectively; Fig 2B, right). We next tested whether simultaneous overexpression of Crk, using UAS-driven mNeonGreen-tagged Crk, rescued effects of knockdown. Crk overexpression alone, even at ∼10-fold endogenous levels (Fig. 2D), had no effect on embryonic viability (99.3%, n=376; Fig. 2C). Simultaneously overexpressing Crk in the *crkS-RNAi* background restored viability to wild-type levels (96.9% viability, n=385; Fig. 2C), supporting the idea that embryonic lethality of Crk knockdown was due to on-target effects. Immunoblotting revealed that this restored total Crk levels (tagged plus endogenous) to 11-15% of normal endogenous levels (Fig 2D). To verify this effect was due to Crk overexpression and not due to titration of GAL4 by the presence of additional UAS-driven elements, we overexpressed either GFP (Valium10 GFP) or a Golgi marker (UASp-GFP.Golgi) in the *crkS-RNAi* background. Neither substantially rescued lethality (9.3% viability (n=387) and 9.0% viability (n=391) respectively; Fig. 2C). We further verified the essential role of Crk in embryonic development using our FRT-flanked allele, by expressing FLP recombinase in the female germline to delete the FRT-flanked *crk* gene, creating females lacking maternally-contributed Crk. When these females were crossed to *crk* heterozygous males, 78% of the embryos died (n=698), consistent with fully penetrant maternal-zygotic lethality and only partial ability of the paternal zygotic gene to rescue viability. Thus, Crk is essential for embryonic viability.

### Loss of Crk disrupts embryonic morphogenesis and leads to formation of multinucleate cells

As a first assessment of Crk’s role in embryonic development, we examined the cuticle secreted by the epidermis to ask whether cell fates were properly determined, major morphogenetic movements like head involution and dorsal closure successfully completed, and whether the epidermis remained intact (Fig. 3A-G). *crkS-RNAi* and *crkW-RNAi* embryos exhibited certain characteristic defects, though at different frequencies (Fig. 3A-H). *crkS-RNAi* embryos had stronger phenotypes, with substantial disruption of epidermal integrity in most embryos. In 25% apparent cell loss occurred, leading to deletion or fusion of denticle belts (Fig. 3B,G, red arrows, H). 20% had holes in the head, ventral or dorsal cuticle (Fig 3C arrow, H), while 11% had more substantial pattern disruption (Fig. 3D arrows, H), and 21% retained only fragments of cuticle (Fig. 3E arrow, H). In embryos in the first three categories, we also noted mild to complete failure to complete germband retraction in 41% (Fig. 3F,G vs A, magenta to black arrows). We suspect the phenotypic spectrum reflects the fact that embryos can inherit one or two shRNA constructs. After *crkW-RNAi* most embryos secreted largely wildtype cuticles (87%, including the 71% that hatched; Fig 3H). However, in 8% deletion or fusion of denticle belts was observed (Fig 3H). 20% of the lethal embryos were also defective in germband retraction. We next generated embryos maternally mutant for Crk, using FLP recombinase to excise the genomic rescue construct in the female germline—half of these were also zygotically mutant. The spectrum of defects was quite similar to those seen in *crkS-RNAi*. Roughly half, potentially those receiving a wildtype gene zygotically, had wildtype cuticles or deletion or fusion of denticle belts (Fig. 3H). The other half had more severe disruptions of the embryonic pattern or only fragments of cuticle (Fig 3H). Defects in head involution, germband retraction and epidermal integrity are all shared with *abl* maternal/zygotic mutants (Grevengoed *et al*., 2001; Rogers *et al*., 2016). The more severe pattern disruptions were similar to those we observed in embryos maternally mutant for the septin protein Peanut (Adam *et al*., 2000), suggesting the possibility that, as in *peanut* mutants, the underlying defects might occur very early, when many cells were lost at or right after cellularization.

**Figure 3.**
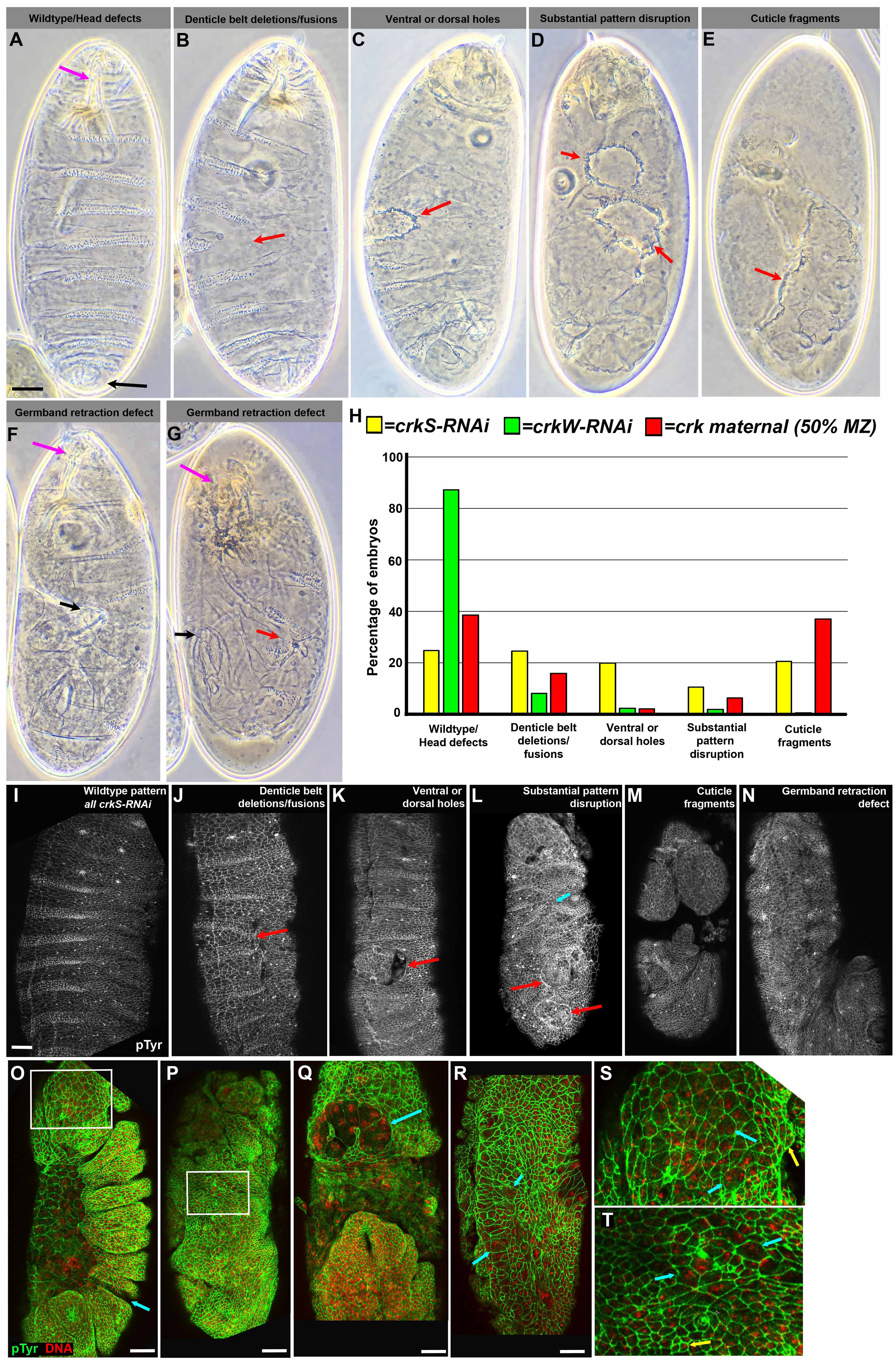
Crk loss disrupts embryonic morphogenesis and leads to the formation of multinucleate cells. A-G. Embryonic cuticles, anterior at top. Range of phenotypes in *crk* maternal-zygotic mutants or after *crkS-RNAi or crkW-RNAi*. A. Wildtype. Note intact, correctly patterned epidermis, and correct completion of germband retraction with head skeleton at anterior (magenta arrow) and spiracles at posterior (black arrow). B. Some mutant or knockdown embryos had deletion or fusion of denticle belts (arrow), though epidermis remained intact. C. Others had holes in anterior, dorsal, or ventral cuticle (arrow). D. In severe cases there were multiple segmental deletions or epidermal holes (arrows). E. In the most severe cases only fragments of cuticle remained (arrow). F. Embryo with a wildtype pattern but in which germband retraction failed, leaving the spiracles (black arrow) curled around the dorsal side. G. Embryo combining denticle belt deletion (red arrow) and germband retraction failure (black arrow). H. Frequencies of defects illustrated. *crk* maternal-zygotic mutants and *crkS-RNAi* lead to a similar range of defects while those caused by *crkW-RNAi* are generally milder. I-N. Stage 15 embryos stained with an antibody to phosphotyrosine (pTyr), revealing cell outlines. Defects in morphogenesis mirror those seen later in the cuticle, including denticle belt fusions (J, arrow) and epidermal holes (K,L red arrows). O-T. Embryos stained for pTyr and a DNA dye. O,P. Stage 13-14. Note pattern disruptions or epidermal holes (arrows). Boxed areas are enlarged in S and T. Q. Stage 11. Note large multinucleate cells in the head (arrow). R. Stage 9. Many large multinucleate cells are seen in the ectoderm (arrows). S,T. Closeups from embryos in O and P, contrasting normal cells (yellow arrows) with larger multinucleate cells (cyan arrows). Scale bars = 30µm.

We extended this analysis by staining embryos with antibodies to phosphotyrosine (pTyr) to outline cells, allowing rapid assessment of morphogenesis. We first examined embryos after morphogenesis should largely be complete, at dorsal closure or later. We observed embryos with a range of defects reflecting those seen in the cuticles (Fig 3I-N). Some were wildtype (Fig 3I), while others had mild disruptions of the epidermal pattern or small holes in the epidermis (Fig 3J,K, arrows). In others morphogenesis was more severely disrupted, with defects in head involution or more severe disruptions in epidermal patterning (Fig. 3L, cyan arrow) or integrity (Fig 3L, red arrows: M). Embryos with germband retraction defects were also observed (Fig 3N). Earlier stage mutants similarly exhibited gaps in the embryonic pattern and holes in the ectoderm (Fig 3O-P). Most striking, we noted frequent examples of multinucleate cells in the developing head (Fig 3O, box enlarged in S; Q) and at a lower frequency in the thorax and abdomen (Fig 3P, box enlarged in T). Once again, these were defects we had previously observed in both *peanut* (Adam *et al*., 2000) and *abl* maternal/zygotic mutants (Grevengoed *et al*., 2003), both of which have defects in syncytial development and/or cellularization. These results prompted us to begin a detailed analysis at these stages.

### Loss of Crk disrupts the cytoskeletal events driving syncytial development

Our data suggested Crk knockdown or loss causes defects that precede the onset of morphogenesis at gastrulation. *Drosophila* begins development as a syncytium with a series of nuclear divisions without cytokinesis, and thus we examined these syncytial divisions to define the earliest role of Crk. The last four rounds of nuclear division occur after nuclei have migrated to the embryo cortex, thus setting the initial membrane and cytoskeletal polarity of the system (Mavrakis *et al*., 2009). The nuclei become increasingly closely packed through these four rounds of mitosis and there is a dynamic cycle of rearrangement of the actin and myosin cytoskeletons that prevents collision of adjacent mitotic spindles, which could lead to aneuploidy. During interphase, actin forms “caps” at the cortex above each nucleus. As spindle assembly begins, these actin caps expand, and when adjacent caps meet, the cytoskeleton is driven inward into transient furrows that surround each mitotic spindle and insulate it from its neighbors (Zhang *et al*., 2018). These pseudocleavage furrows recede as mitosis is completed and caps re-form, beginning a new cycle. In mutants in which furrow formation is blocked or reduced (e.g. Sullivan *et al*., 1993; Afshar *et al*., 2000; Zallen *et al*., 2002; Webb *et al*., 2009), spindles often collide, leading to DNA damage and/or aneuploidy. This triggers a Chk2-mediated response by which aneuploid nuclei are removed from the embryo surface by “dropping” into the interior of the egg during the next mitotic cycle (Takada *et al*., 2003). At the end of nuclear cycle 13, furrows form around each nucleus and pull membranes downward, enclosing each nucleus in a cell, in a process called cellularization. Actin rings stabilize yolk channels at the bottom of each new cell, connecting it to the underlying yolk.

In wildtype the dynamic cycle of actin rearrangements ensures most syncytial mitoses occur without errors. Nuclei are well spaced at cycle 10, not all caps encounter a neighbor, and furrows are relatively shallow. Spindles assemble, and chromosomes segregate without errors (Fig. 4A). As each cycle proceeds, nuclear density increases (Fig. 4C,E,G,J)—however, pseudocleavage furrows generally succeed in maintaining spindle separation. In wildtype, occasional defects in spindle separation and chromosome segregation occur in later cycles (1/8 and 4/19 cycle 11 and 12 embryos had 1-2 spindle collisions per field of view; Fig. 4P). Defective nuclei are removed into the yolk in the next cycle. This affects 2-3% of nuclei in wildtype (Sullivan *et al*., 1993). Centrosomes remain behind and continue to organize the actin cytoskeleton into smaller caps and furrows (Fig. 4G, arrow). Nuclear loss tends to affect small groups of neighboring nuclei. In wildtype, increasing numbers of cycle 11 or 12 embryos had small patches of nuclear loss (4/8 and 13/19 at cycles 11 and 12, relative to 1/10 in cycle 10), but there were only 1-4 patches per embryo and these included only a relatively small number of nuclei (e.g. Fig 4G, arrow).

**Figure 4.**
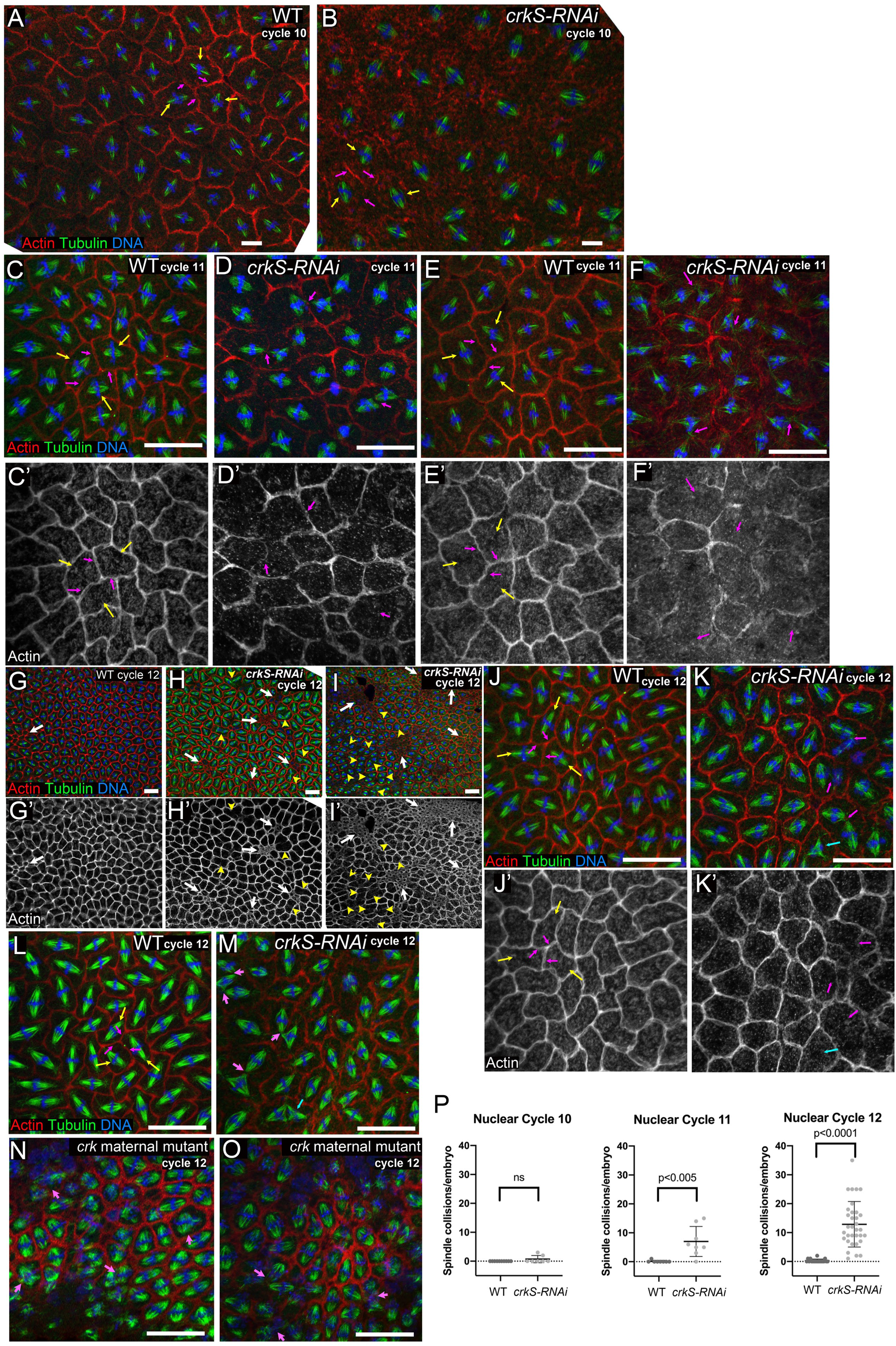
Crk knockdown disrupts the cytoskeletal events driving syncytial development. A-O. Embryos in metaphase to anaphase of indicated syncytial nuclear cycle, stained to visualize F-actin, tubulin, and DNA. A,B. Nuclear cycle 10. A. Wildtype. Each spindle and its associated chromosomes (yellow arrows) are surrounded by a pseudocleavage furrow (magenta arrows), separating nuclear compartments and preventing spindle collisions. B. *crkS-RNAi.* Spindles remain separated from one another (yellow arrows), though pseudocleavage furrows were often less apparent (magenta arrows). C-F. Nuclear cycle 11. C,E. Wildtype. Spindles remain well separated (yellow arrows) and pseudocleavage furrows continue to separate nuclear compartments (magenta arrows). D,F. *crkS-RNAi.* Most embryos have multiple spindle collisions (magenta arrows). These usually occur in places where pseudocleavage furrows are discontinuous or absent. G-O. Nuclear cycle 12. G. Most wildtype embryos have a small amount of nuclear loss, revealed by small actin rings, usually only involving a few nuclei (arrow). Spindle collisions remain very rare. H,I. *crkS-RNAi.* Most embryos have multiple regions of nuclear loss (white arrows), and in severe cases these occupy a significant fraction of the embryo surface (I, arrows). Spindle collisions (yellow arrowheads) occur in regions adjacent to and distant from regions of nuclear loss. J,L. Wildtype. Spindles (yellow arrows) remain separated by pseudocleavage furrows (magenta arrows). K,M. *crkS-RNAi*. All embryos have multiple spindle collisions (magenta arrows), sometimes leading to tripolar spindles (cyan arrows). Spindle collisions occur in regions where pseudocleavage furrows are weak or absent. N,O. *crk* maternal mutants exhibiting spindle collisions (arrows) and nuclear loss. P. Quantification, number of spindle collisions per embryo in wildtype and *crkS-RNAi* in nuclear cycles 10-12. Spindle collisions are elevated in cycle 11 and highly elevated in cycle 12. Scale bars=20µm.

We first examined Crk localization during syncytial divisions, using our mNeonGreen-tagged construct. Strikingly, Crk is enriched in both actin caps (Fig. 1H) and metaphase furrows (Fig. 1I), consistent with the possibility that Crk plays a role in cytoskeletal regulation. We next examined the effect of Crk knockdown. In *crkS-RNAi* embryos, nuclear cycle 10 proceeded relatively normally (Fig. 4B), though pseudocleavage furrows appeared less continuous (see below), and 2/7 embryos observed had individual examples of spindles that had collided (quantified in Fig. 4P). However, by nuclear cycle 11, spindle defects became more frequent: in 7/8 embryos examined neighboring spindles collided (Fig. 4D,F, magenta arrows), often in places where pseudocleavage furrows were reduced or absent. Embryos often had multiple examples in a single field of view (quantified in Fig. 4P). At cycle 11, the frequency of embryos with regions of nuclear loss remained similar to that in wildtype (4/8 embryos observed). However, by nuclear cycle 12 all *crkS-RNAi* embryos (33/33) had multiple spindle collisions, and in the most severely affected these included a substantial fraction of the spindles in the field of view (Fig 4H,I, arrowheads; K,M magenta arrows; quantified in Fig. 4P). The incidence of nuclear loss increased (91% affected versus 68% in wildtype), and the fraction of nuclei involved was much higher, with 71% having multiple groups of nuclei lost, including embryos in which this affected much of the embryo surface (Fig. 4H-I, white arrows). We think the extensive nuclear loss likely reflects the frequency of spindle collisions in the previous cycle. Spindle collisions and nuclear loss were also observed in *crk* maternal mutants generated using our FRT flanked genomic constructs (Fig. 4N,O). These data reveal Crk plays an important role in regulating the intricate cytoskeletal rearrangements of syncytial development.

At the end of the syncytial stage, cellularization forms the first cells (Lee and Harris, 2014; Schmidt and Grosshans, 2018). An actomyosin ring assembles at the cortex above each nucleus—the rest of the cortex is covered with actin-rich microvilli. Cofilin, Anillin, Septins, Rho-driven myosin contractility, regulated actin assembly involving the formin Diaphanous, and regulation of membrane dynamics all play roles (e.g., Grosshans *et al*., 2005; Padash Barmchi *et al*., 2005; Figard *et al*., 2016; Xue and Sokac, 2016), driving the movement of these rings inward, enclosing each nucleus in a plasma membrane (Fig. 5A-D, I). Both actin and myosin localize to the leading edge during this event (Fig. 5A-D, cyan arrows). At the end of cellularization, the actomyosin rings constrict to partially close off each cell, but this halts before completion, leaving an actin-lined “yolk channel” at the basal end of each cell (Fig. 5K). Abl plays a role in this process (Grevengoed *et al*., 2003), and many genes involved in syncytial divisions also play a role in cellularization. We thus examined cellularization after *crk* RNAi.

**Figure 5.**
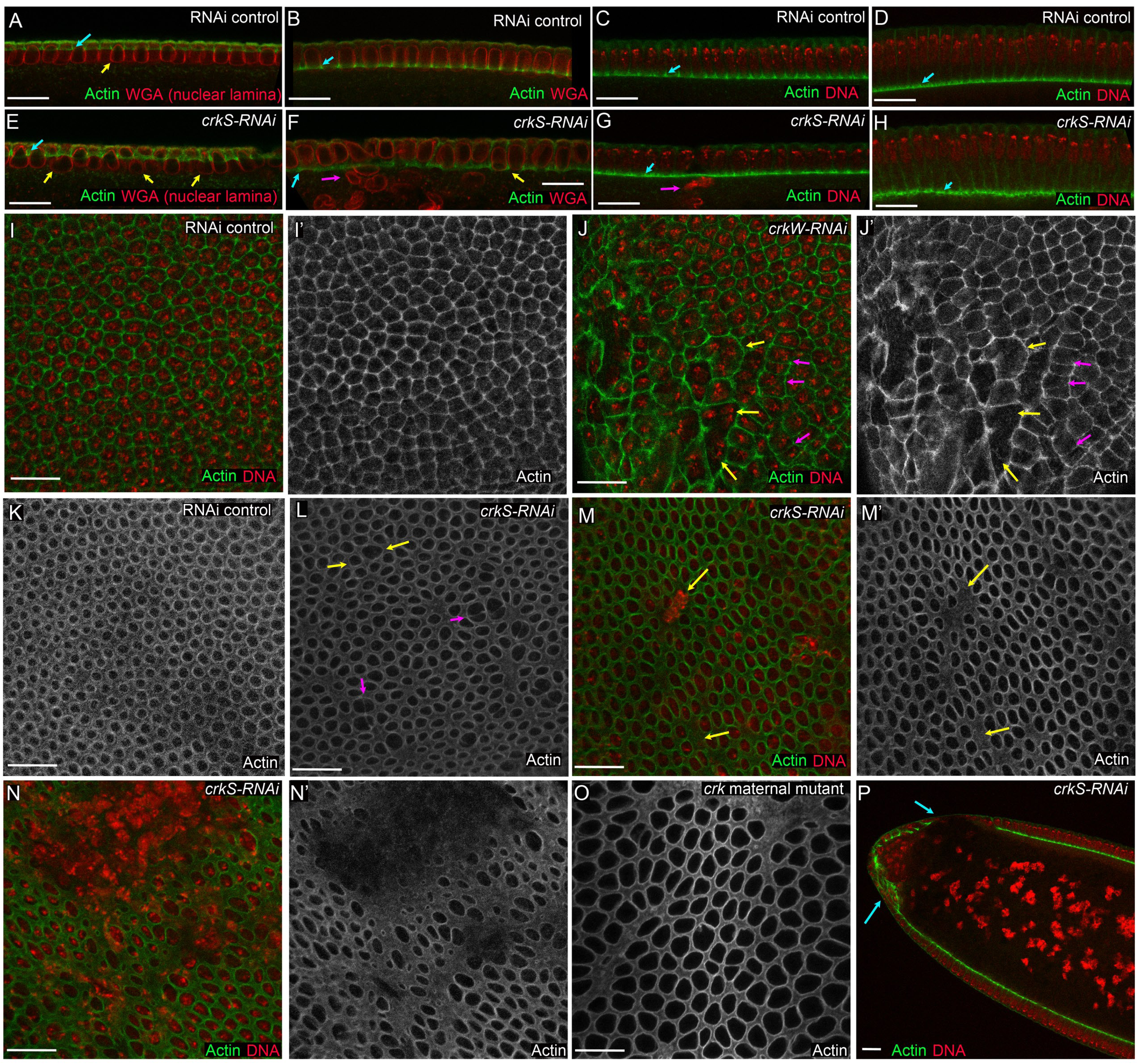
Crk knockdown leads to defects in cellularization. Cellularizing control, *crkS-RNAi* and *crk* maternal mutant embryos stained with phalloidin to visualize F-actin and either WGA to visualize the nuclear lamina or a DNA dye. A-H. Cross section views during early (A,E), mid- (B vs, F,C vs G,) and late cellularization (D,H). A-D. In controls, as in wildtype, actin localizes to cellularization front (cyan arrows). moving basally as a ring around each nucleus and then partially closing, leaving the yolk channel. Apical actin-rich microvilli gradually disappear during this process. Nuclei are uniform in shape (e.g. A, yellow arrow) and positioned in a regular row. E-H. *crkS-RNAi*. Actin can assemble at the front and move basally (cyan arrows). However, nuclear shape is irregular (E, F yellow arrows), possibly reflecting premature ring closure. Some nuclei reside below the cortex (F,G, magenta arrows), likely a result of nuclear loss in syncytial cycles. I,J. *en face* views, mid-cellularization. I. Wildtype. Actin rings are uniform in size and actin intensity and enclose single nuclei. J. *crkS-RNAi*. Note multinucleate actin compartments (yellow arrows) and examples of weak actin localization to a subset of compartment boundaries (magenta arrows). K-O. Late cellularization at level of forming yolk channels. K. Wildtype. Rings partially close, leaving actin-lined yolk channels of uniform size. L,M. Examples of milder defects after *crkS-RNAi*. Yolk channels are uneven in size (L, yellow arrows) and sometimes two channels are separated only by a weak actin border (L, magenta arrows). Regions without yolk channels are also seen (M, yellow arrows), some of which coincide with places where nuclei lie below the cortex. N. Severe example of *crkS-RNAi* defects with a large region of nuclear loss. O. *crk* maternal mutant exhibiting similar variability in yolk channel size. P. Cross section, *crkS-RNAi* embryo at gastrulation onset. Many embryos have severe cellularization defects in the presumptive head (arrows). Scale bars=20µm.

Several defects were readily apparent in most *crkS-*RNAi embryos. While in controls nuclei form a uniform row of similar shape (Fig. 5A-C), in *crkS-*RNAi embryos nuclei were seen below this row (Fig. 5F,G, magenta arrows), likely representing nuclei removed from the surface in the previous syncytial division. Nuclear shape was less uniform than in controls (Fig. 5E,F yellow arrows vs A,B), a phenotype that can reflect premature closure of the actomyosin ring—the so-called bottleneck phenotype (Schejter and Wieschaus, 1993). Viewed *en face*, actin rings in wildtype embryos are largely uniform in shape and intensity, and each encloses a single nucleus (Fig. 5I). In contrast, after *crk* RNAi, actin rings were less uniform, the actin staining at the border between two nuclear compartments was sometimes less intense (Fig. 5J, magenta arrows), and in other cases it was absent, leaving multiple nuclei in a single membrane compartment (Fig. 5J, yellow arrows). This disruption of nuclear compartmentalization is a likely mechanism for the formation of multinucleate cells we observed above. We also observed striking defects in yolk channels in Crk-depleted embryos. While these are uniform in size and shape in wildtype (Fig. 5K), they varied dramatically in size and shape following *crkS-RNAi* (Fig. 5L-N). Some channels were abnormally enlarged (Fig 5L, yellow arrows), and others appeared to be separated by only a weak actin border (Fig. 5L, magenta arrows). Other regions were devoid of yolk channels altogether—these often overlaid nuclei that had been removed from the cortex (Fig. 5M, arrows). In the most severe cases, large regions of the embryo were affected (Fig. 5N), as we had observed at the syncytial stages—these were prominent in the presumptive head as gastrulation began (Fig. 5P, arrows). Defects in cellularization leading to variable size yolk channels were also observed in *crk* maternal mutants generated using our FRT flanked genomic construct (Fig. 5O). Intriguingly, live imaging revealed the ability of embryos to “repair” the regions devoid of nuclei (Suppl. Movie 1), helping explain the embryos with pattern deletions we observed at later stages. Together, these data reveal Crk is important for proper regulation of the dynamic behavior of the actomyosin cytoskeleton during syncytial divisions and cellularization.

### Crk is important for actin cap expansion and the resulting establishment of pseudocleavage furrows

In some mutants with similar phenotypes, the primary defect is in proper formation of pseudocleavage furrows, which form as expanding actomyosin caps collide (Zhang *et al*., 2018). We thus focused our attention on this event, measuring maximum furrow depth at metaphase. We drew lines across furrows separating neighboring nuclei (e.g., Fig. 6 A-F), reconstructed z-axis projections from z-stacks, and used these to measure furrow depth. In wildtype, pseudocleavage furrows first form during cycle 10, and then become sequentially deeper in the subsequent cycles. We compared furrow depths in wildtype to those in *crkS-RNAi* embryos, choosing pairs of nuclei without spindle collisions to avoid events biased by earlier furrow failure. As our *en face* views had suggested (Fig. 6 A vs B), pseudocleavage furrows invagination was significantly shallower at cycle 10 (Fig. 6G, mean 1.96 µm in wildtype versus 1.09µm in *crkS-RNAi*). This defect remained significant at cycles 11 (Fig. 6H, mean 2.42 µm in wildtype versus 1.89µm in *crkS-RNAi*) and 12 (Fig. 6I, mean 2.69 µm in wildtype versus 2.06 µm in *crkS-RNAi*). While remaining furrows were similar in depth at nuclear cycle 13 (Fig. 6J; mean 3.18 µm in wildtype versus 3.26 µm in *crkS-RNAi*), by that stage spindle collisions were so frequent in *crkS-RNAi* embryos that we could only measure furrows in the least affected embryos.

**Figure 6.**
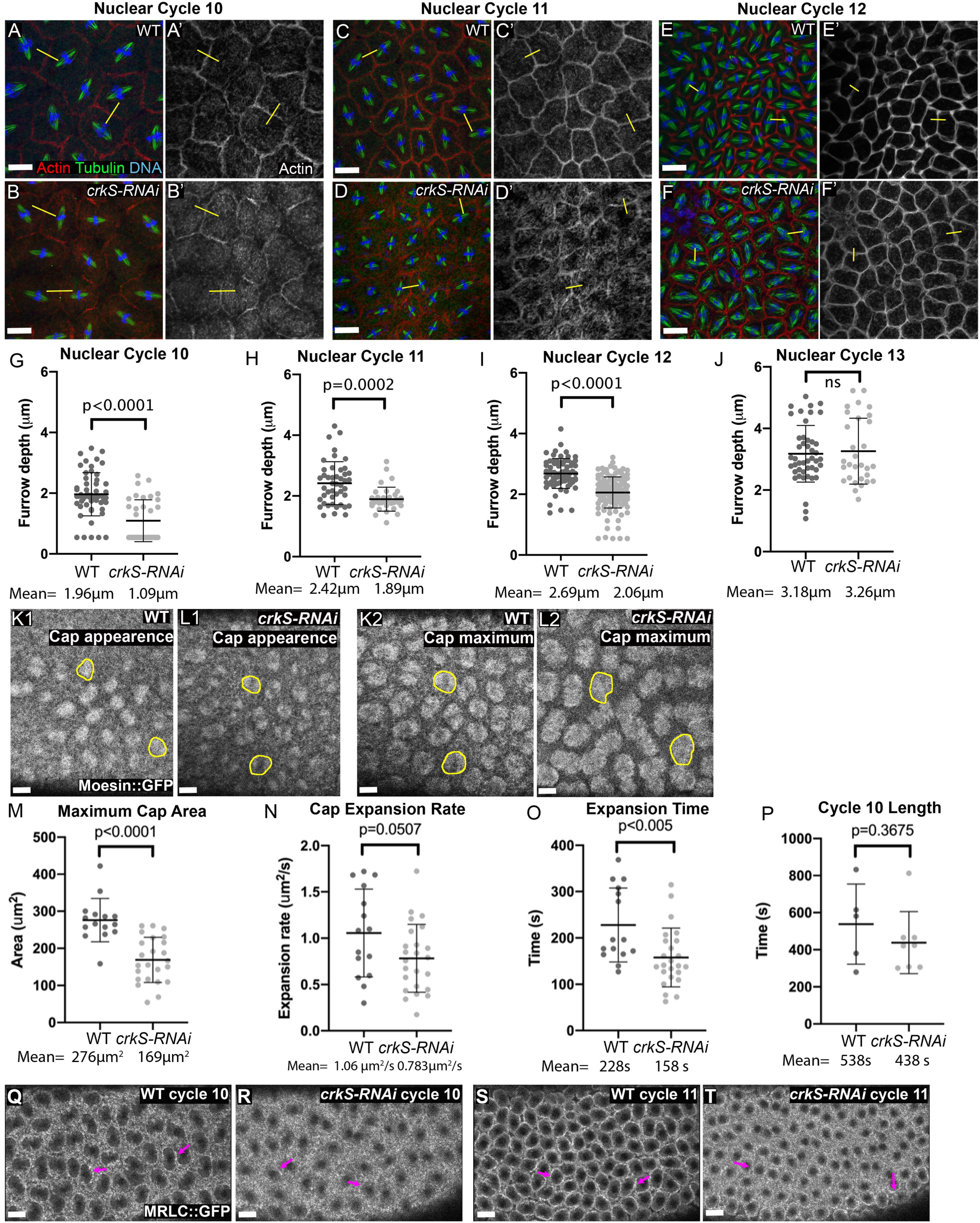
Crk knockdown reduces pseudocleavage furrows depth and cap expansion. A-F. Representative images, wildtype and *crkS-RNAi* embryos at nuclear cycles 10, 11 and 12. Embryos fixed and stained to visualize actin, tubulin and DNA, selected for those in metaphase to anaphase, and binned by nuclear density into the respective cycles. Furrows were selected, avoiding those involved in apparent mitotic spindle collisions. Lines were drawn perpendicular to the spindle (examples in yellow), z-axis projections were made, and metaphase furrow depth was measured (see Methods for details). G-J. Quantification of furrow depth. Furrows were significantly less deep in nuclear cycles 10-12 in *crkS-RNAi* embryos. K-L. Movie stills, illustrating examples of wildtype embryos at first appearance of actin caps and at the point when caps were maximally expanded—outlines are examples of caps measured for the analysis. Full movie=Suppl. Movie 2. M-P. Quantification of cap dynamics (see Methods for definitions). Q-T. Stills from movies of wildtype and *crkS-RNAi* embryos expressing MRLC::GFP, visualized at the peak of cap expansion in nuclear cycles 10 and 11. Note reduction in myosin accumulation at cap margins after *crkS-RNAi*. Full movie = Suppl. Movie 3. Scale bars=20µm.

We also examined cap expansion directly, by using imaging of GFP fused to the F-actin binding domain of Moesin (sGMCA; Edwards *et al*., 1997) to visualize the actin cytoskeleton in living embryos. Our movies suggested cap expansion was less extensive in mutants (Suppl. Movie 2); this was particularly striking in the early cycles in *crkS-RNAi* embryos. To quantify cap expansion, we focused on cycle 10, when increased cap spacing allowed more accurate measurement of individual cap area. We measured cap area both when caps first appeared at the surface of the embryo (Fig. 6K1,L1) and again when they reached maximal expansion (Fig. 6K2,L2; see Methods for details). While starting cap area was similar between wildtype and *crkS-RNAi* embryos, maximum cap area was significantly reduced after *crkS-RNAi* (Fig 6M). We calculated the cap expansion rate by plotting the change in cap area over time and found it was reduced following *crkS-RNAi* (Fig 6N), partially due to premature termination of cap expansion (Fig 6O). However, overall length of nuclear cycle 10 (measured from first cap appearance to the next cap appearance) was not significantly changed (Fig. 6P).

As actin caps collide, myosin becomes enriched at the cap borders and furrows (Zhang *et al*., 2018). We visualized myosin live through the syncytial cycles using a GFP-tagged form of Myosin Regulatory Light Chain (MRLC=Sqh::GFP; Suppl. Movie 3). Stills from the end of cap expansion during nuclear cycles 10 and 11 are shown in Fig. 6Q-T. In wildtype myosin is clearly enriched at cap margins at both nuclear cycles (Fig. 6Q,S, arrows). In contrast, myosin enrichment at cap margins appeared substantially reduced in *crkS-RNAi* embryos (Fig. 6R,T, arrows). Together, these data suggest the defects in syncytial divisions in *crkS-RNAi* embryos result from defects in actin dynamics, with reduced cap expansion leading to shorter pseudocleavage furrows and subsequent spindle collisions and nuclear loss.

### Crk knockdown reduces levels of Arp2/3 and its activator SCAR in actin caps and pseudocleavage furrows

The reduced expansion of actin caps and impaired establishment of pseudocleavage furrows, suggested the possibility that the localization and/or activity of actin cytoskeletal regulators was altered by loss of Crk. The Arp2/3 complex (Stevenson *et al*., 2002) and its upstream activator SCAR (Zallen *et al*., 2002) are both required for actin cap expansion and pseudocleavage furrow formation during syncytial development. Since Crk regulates Arp2/3-dependent actin remodeling in other cellular contexts (e.g., myoblast fusion, phagocytosis, and immune synapse maturation), we hypothesized localization and/or activity of the Arp2/3 complex or SCAR might be altered by loss of Crk. We tested this hypothesis using quantitative imaging of Arp3 and SCAR localization to actin caps and pseudocleavage furrows. To directly compare fluorescence intensity between genotypes, we stained *crkS-RNAi* embryos together with embryos marked with Histone::RFP as internal, wild-type controls. Compared to controls, SCAR intensity in both actin caps (Fig. 7B compared to A) and pseudocleavage furrows (Fig. 7D compared to C) was significantly decreased following *crkS-RNAi* (Fig. 7F,H). Arp3 localization to both actin caps (Fig. 7J compared to I) and pseudocleavage furrows (Fig. 7L compared to K) was also significantly reduced by *crkS-RNAi* (Fig. 7N,P). Quantification of phalloidin intensity revealed F-actin levels were, on average, significantly reduced in both actin caps (Fig. 7B and J compared to A and I, respectively; quantified in E,M) and pseudocleavage furrows (Fig. 7D and L compared to C and K, respectively; quantified in G, O). Taken together, these data suggest the altered actin cap dynamics and impaired pseudocleavage furrow formation observed after *crkS-RNAi* result from altered localization and thus activity of the Arp2/3 complex and its upstream activator, SCAR.

**Figure 7.**
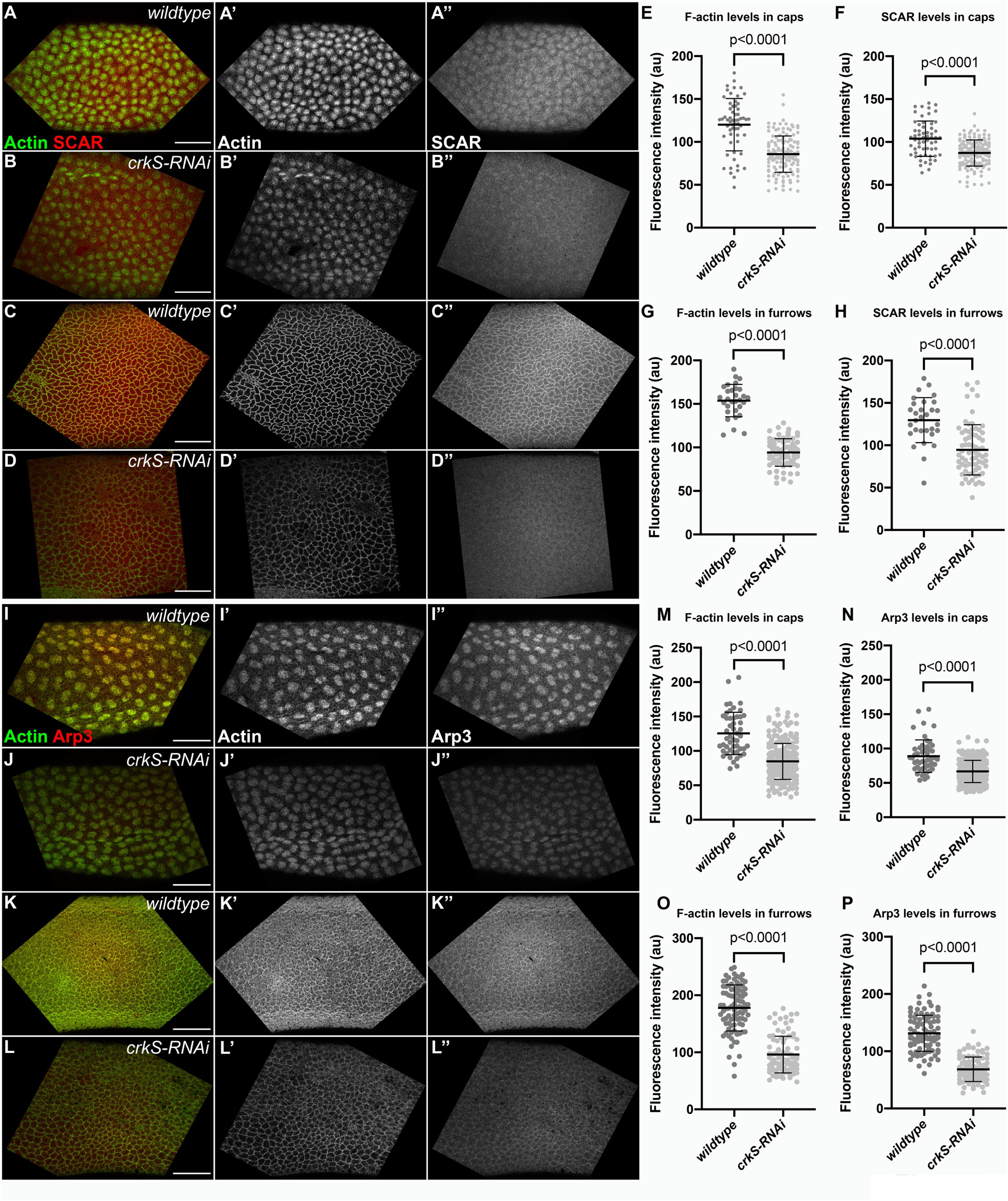
Crk knockdown leads to reduced localization of the Arp2/3 activator SCAR, Arp3, and F-actin to caps and pseudocleavage furrows. A-D. Representative images, wildtype and *crkS-RNAi* embryos stained with phalloidin (visualizing F-actin) and an antibody against SCAR. E-H. Quantification, fluorescence intensity of F-actin or SCAR in actin caps (E,F) and pseudocleavage furrows (G,H). In wildtype, SCAR localizes to both actin caps (A-A’) and pseudocleavage furrows (C- C’). SCAR still localizes to actin caps (B-B”) and pseudocleavage furrows (D-D”) following Crk knockdown, but levels of F-actin and SCAR in both structures is significantly reduced after *crkS-RNAi* (E-H). I-L. Representative images, wildtype and *crkS-RNAi* embryos stained with phalloidin and an antibody against Arp3. M-P. Quantification, fluorescence intensity of F-actin and Arp3 in actin caps (M,N) and pseudocleavage furrows (O,P). Arp3 also localizes to actin caps and pseudocleavage furrows in both wildtype (I-I’’, K-K’’) and *crkS-RNAi* (J-J’’, L-L’’) embryos. Like SCAR, Arp3 localization to both actin caps (N) and pseudocleavage furrows (P) is significantly reduced after *crkS-RNAi*. F-actin levels in actin caps (M) and pseudocleavage furrows (O) is also significantly reduced after *crkS-RNAi*. In all graphs, mean=black bar; error bars=standard deviation; n=6 wildtype and 14 crkS-RNAi embryos in E,F; n=3 wildtype and 7 *crkS-RNAi* embryos in G,H; n=5 wildtype and 22 *crkS-RNAi* embryos in M,N; n=9 wildtype and 9 *crkS-RNAi* embryos in O,P. All scale bars=50µm.

### Loss of Crk leads to defects in axon patterning in the central nervous system similar to those seen in *abl* mutants

Based on Crk’s multiple roles in mammalian cells, we suspected Crk likely played additional roles in *Drosophila* embryogenesis. We initiated work on Crk because we suspected it was a regulator or effector of Abl kinase, as the PXXP motif within Abl that can bind Crk is essential for Abl function (Rogers *et al*., 2016). Abl plays many roles in embryonic morphogenesis, but the best characterized are its roles in the central nervous system (CNS), where it regulates axon outgrowth and pathfinding (e.g. Gertler *et al*., 1989; Wills *et al*., 1999a; Wills *et al*., 1999b; Bashaw *et al*., 2000; Liebl *et al*., 2000; Crowner *et al*., 2003; Forsthoefel *et al*., 2005). We first examined whether Crk was expressed in the CNS, using our mNeonGreen-tagged rescue construct. Crk localized to the axon tracks of the CNS (Fig 8A, cyan arrow), as well as to midline glia (Fig 8A, magenta arrow), suggesting Crk may play a role in axon patterning. We thus examined the nervous systems of Crk knockdown embryos, using both our strong and weak shRNA lines. We visualized axons using antibodies to BP102, which labels all CNS axons (Seeger *et al*., 1993), and Fasciclin II (FasII), an adhesion molecule expressed on a subset of CNS axons (Grenningloh *et al*., 1991).

**Figure 8.**
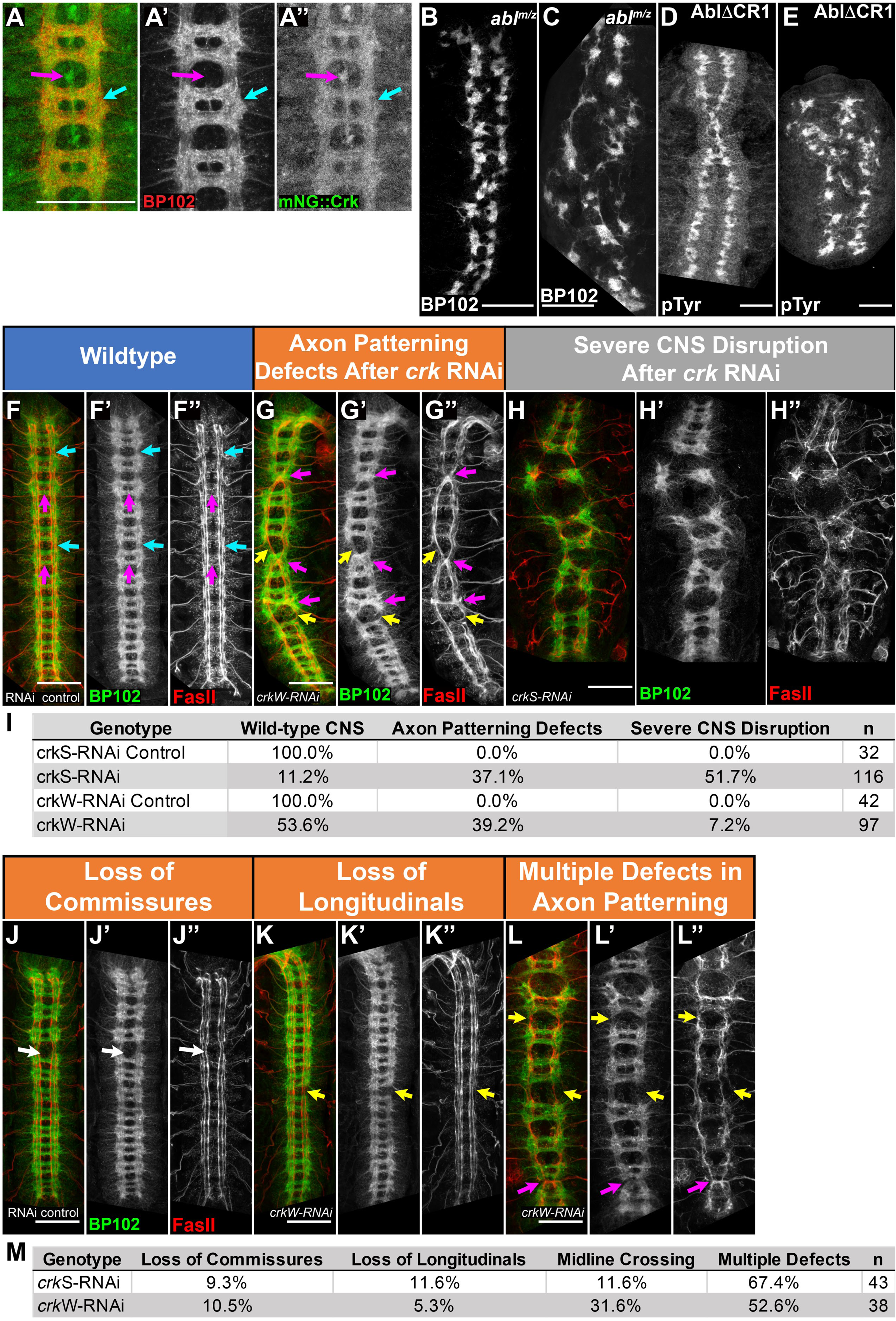
Crk is required for proper patterning of the embryonic central nervous system. A-H, J-L. Stage 15 embryos, antigens and genotypes indicated. I,M. Quantification of penetrance of phenotypes depicted. A. mNG::Crk expressed at endogenous levels. Crk is enriched in CNS axons (cyan arrow) and midline glia (magenta arrow), suggesting Crk may play a role in axon pathfinding or function. B-C. Representative images, CNS phenotypes, *abl* maternal and zygotic null mutants. D-E. CNS phenotypes, *abl* mutants expressing a form of Abl (AblΔCR1) that cannot bind to PXXP-dependent partners. Both can lead to loss of commissures (B,D) or severe disruption of CNS patterning (C,E). F-I. Representative images, range of CNS phenotypes in *crkS-RNAi* and *crkW-RNAi* embryos and quantification of their prevalence (I). In wildtype, axons form a stereotypical ladder-like pattern with longitudinal axon bundles running anterior to posterior (cyan arrows, F-F’’) and commissural axon bundles crossing the midline at defined points in each body segment (magenta arrows, F-F’’). FasII-positive axons run in three parallel bundles on either side of the midline and do not normally cross the midline (F’’). Crk knockdown results in a range of CNS phenotypes, including milder axon patterning defects and severe disruption of CNS patterning (G-H’’,I). J-M. Representative images, different axon patterning defects observed after Crk knockdown and quantification of their prevalence. These include aberrant midline crossing (magenta arrows, G-G’’, L-L”), loss of longitudinal axons (yellow arrows, G-G’’, K-K”, L-L”), and loss of commissural axon bundles (white arrow, J-J’’). All scale bars=50µm.

In wildtype, CNS axons form a ladder-like array (Fig. 8F), with axons extending longitudinally on either side of the midline (Fig 8F, cyan arrows) and crossing the midline at two commissures per body segment (Fig 8F, magenta arrows). FasII-positive axons follow three parallel longitudinal tracks on each side of the midline, and do not normally cross the midline (Fig. 8F”). Embryos maternally and zygotically mutant for null alleles of *abl* exhibit a range of CNS defects (Rogers *et al*., 2016). Roughly half have moderate defects in axon patterning, with frequent loss of commissures (Fig 8B). The other half have more severe defects in axon patterning (Fig. 8C). Embryos mutant for *abl^ΔCR1^,* in which the PXXP motif is deleted, exhibit a similar range of defects (Fig 8D,E), though the penetrance is less complete—22% are wildtype while 70% have commissural defects (Rogers *et al*., 2016). We thus compared these defects to the effects of knocking down Crk.

*crkS-RNAi* led to substantial and highly penetrant defects in axon pattern (Fig 8F-H; quantified in I). In 52% of embryos, there was severe disruption of nervous system patterning (e.g., Fig. 8H). 37% of embryos had milder defects in axon patterning (Fig 8G), with inappropriate midline crossing of FasII-positive axons (Fig. 8G, L magenta arrows), as well as loss of commissural (Fig 8J, arrow) or longitudinal (Fig. 8G,K,L yellow arrows) axons in some segments (quantified in Fig 8M). 11% of embryos had a normal CNS, consistent with the embryonic viability data after strong knockdown. This range of defects was similar to what we previously observed in *abl* maternal/zygotic mutants or after mutation of Abl’s PXXP motif (Fig 8B-E; (Grevengoed *et al*., 2001; Rogers *et al*., 2016), though total loss of commissures was less frequent after Crk knockdown. *crkW-RNAi* produced weaker and less penetrant defects (quantified in Fig. 8I,M). Only 7% of embryos had severe CNS disruption, once again consistent with the reduced embryonic lethality. However, 39% had milder but still significant CNS defects, once again ranging from loss of longitudinal or commissural axons in some segments to inappropriate midline crossing of FasII-positive axons.

Because *crkS-RNAi* leads to major defects in cellularization, including cell loss at gastrulation onset, we suspected many of the severe CNS defects might be a secondary consequence of these earlier defects, with loss of ectodermal cells that should have become neuroblasts. However, *crkW-RNAi* results in fewer major morphogenesis defects (Fig. 3), suggesting some of the milder defects we observed might be due to effects of Crk knockdown in the CNS itself. To assess this, we examined epidermal integrity and CNS patterning in the same embryos, using an antibody to phosphotyrosine (pTyr) to outline epidermal cells and antibodies to BP102 and FasII to examine the underlying nervous system (Suppl. Fig. 3). As we suspected, most embryos with strong defects in the CNS also had strong defects in the overlying epidermis, ranging from epidermal holes to strong disruptions to the epidermal pattern (Suppl. Fig. 3A, B arrows), consistent with early cell loss. In other cases, more subtle epidermal defects, such as denticle belt fusions, also correlated with underlying CNS defects (Suppl. Fig. 3C, arrow). However, we also observed mild CNS defects in some embryos in which the overlying epidermis was intact Suppl. Fig. 3D,E arrows), keeping open the possibility that Crk also plays important CNS-intrinsic roles.

### Crk is not essential for the patterns of expression of the guidance cues Slit and Robo, but may play more subtle roles in Slit/Robo signaling

A variety of axon guidance cues shape the pattern of the CNS. Among these is Slit-Robo signaling. The Robo receptor is expressed on a subset of CNS axons (Fig. 9A, cyan arrows). Slit is expressed by midline glial cells (Fig 9D, magenta arrows), and acts as a repulsive ligand for the Robo receptor, so that axons expressing Robo strictly follow longitudinal axon pathways (Fig. 9A, cyan arrows) and do not cross the midline (Fig. 9A, magenta arrows; Blockus and Chedotal, 2016). Slit/Robo signaling requires Abl function (Bashaw *et al*., 2000). The midline crossing defects seen following Crk knockdown could reflect defects in Slit or Robo expression or function. We assessed this by examining expression patterns of both proteins. In wildtype, Robo is expressed in a subset of axons that all follow the longitudinal axon tracks (Fig. 9A, cyan arrows), and is not expressed on axons that cross the midline (Fig. 9A, magenta arrows). Slit is expressed in glial cells along the midline (Fig. 9D, magenta arrows), where the commissural axons cross. We examined Slit and Robo expression patterns in a subset of Crk knockdown embryos in which the CNS was not totally disrupted. Most Crk knockdown embryos retained clear enrichment of Robo expression in the longitudinal tracts (Fig 9B,C, cyan arrows), though these were often displaced laterally. The fact that most Robo-positive axons remained restricted to longitudinal tracks suggested midline repulsion was not completely inactivated. However, while some axons crossing the midline were Robo-negative (Fig. 9B,C, magenta arrows), a subset of Robo-expressing axons did cross the midline (Fig. 9C, green arrows), suggesting Slit/Robo signaling may be impaired following Crk loss. In Crk knockdown embryos in which the nervous system was only mildly disrupted, Slit expression remained confined to the midline glia (Fig. 9E, magenta arrows). However, in embryos with more severely disrupted nervous systems, Slit-expressing cells were sometimes displaced (Fig. 9F, yellow arrows). Thus, Crk does not appear to be required for the basic expression pattern of Robo or Slit, and its knockdown does not fully abrogate midline repulsion. However, the severity of the CNS disruptions observed made it impossible to definitively determine whether Crk plays more subtle roles in Slit-Robo signaling. Together, this work suggests Crk may play two roles in CNS patterning: one indirect, via maintaining the survival of neural precursors just following cellularization, and one CNS intrinsic in axon patterning. However, this latter role remains speculative.

**Figure 9.**
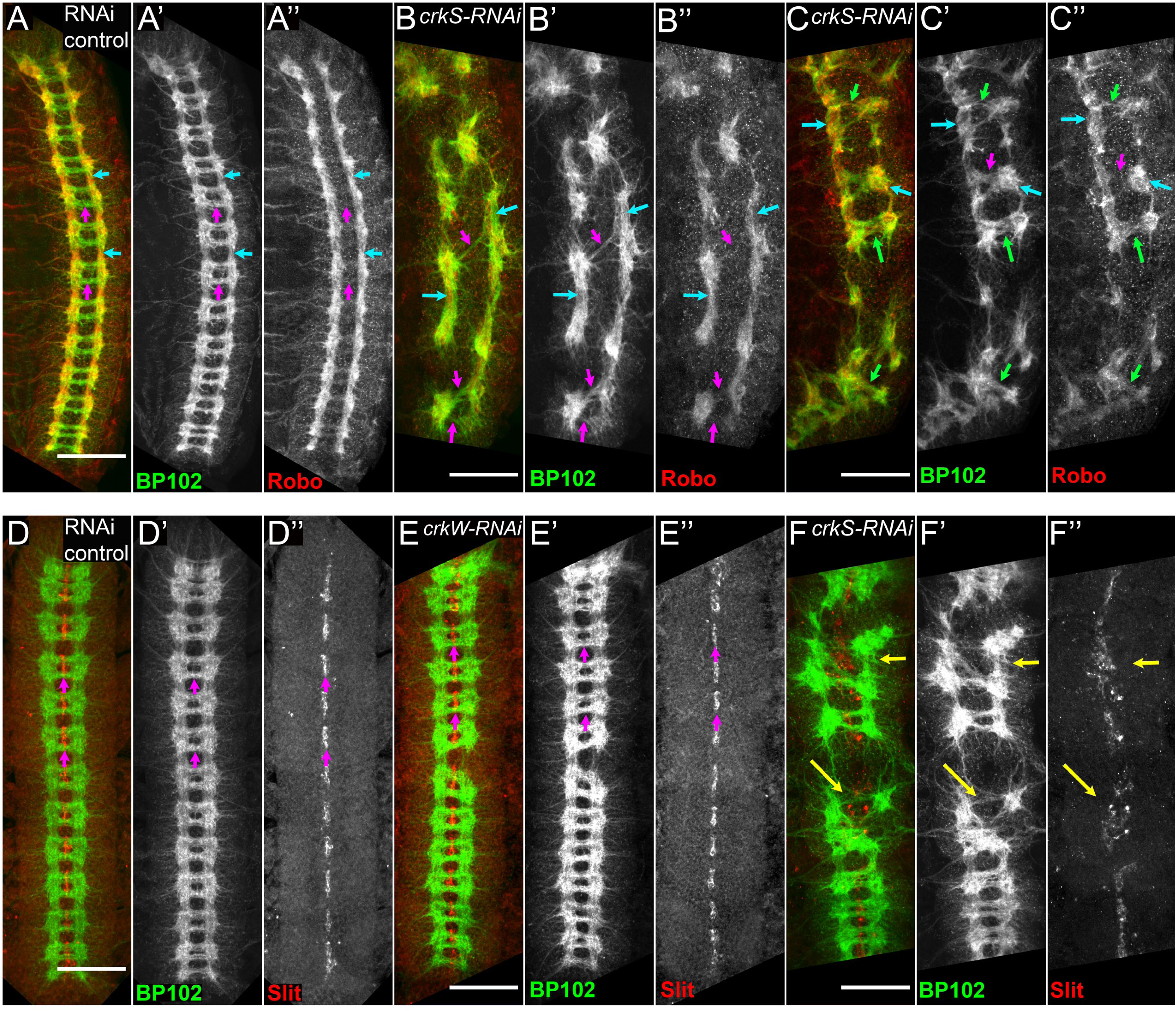
Crk is not required for proper expression of Robo and Slit but may play a more subtle role in Slit/Robo signaling. A-F. Stage 15 embryos, antigens and genotypes indicated. A-C. Representative images of Robo localization in control and Crk knockdown embryos. A. In wild-type, Robo is strongly enriched in longitudinal axon bundles (A-A”, cyan arrows) and excluded from commissural axon bundles crossing the midline (A-A”, magenta arrows). B,C. Following Crk knockdown, Robo expression is still largely restricted to longitudinal axon tracks (B-C’’, cyan arrows). In most cases, midline-crossing axon bundles are Robo-negative (B-C’’, magenta arrows); however, a subset of Robo-positive axons appear to cross the midline in Crk knockdown embryos (C-C’’, green arrows), suggesting Slit/Robo signaling may be impaired following Crk loss. D-F. Representative images, Slit localization in control and Crk knockdown embryos. D. In wildtype, Slit expression is confined to the midline glia along the ventral midline (magenta arrows). E,F. In Crk knockdown embryos, Slit expression remains confined to midline glia, generally forming a stereotypical line along the ventral midline (E=E”’, magenta arrows). However, Crk knockdown embryos with more severely disrupted CNS patterning defects can have mispositioned Slit-expressing midline glia (F-F”, yellow arrows). Scale bars=50µm.

### Crk is required for timely wound healing in the embryonic epidermis

Another embryonic process requiring coordinated regulation of adhesion and actin dynamics is embryonic wound healing (e.g. Martin and Lewis, 1992; Wood *et al*., 2002; Abreu-Blanco *et al*., 2012; Fernandez-Gonzalez and Zallen, 2013). Embryonic wounds are repaired through the collective movements of the cells around the wound. Upon wounding, the cells immediately adjacent to the wound polarize their actin cytoskeleton and assemble a supracellular actin cable that coordinates cell movements. This process is tightly coordinated with dynamic changes in cell-cell adhesion (reviewed in Hunter and Fernandez-Gonzalez, 2017). Abl is required for efficient wound closure, via effects on the organization of filamentous actin and the reorganization of cell-cell junctions at the wound margin (Zulueta-Coarasa *et al*., 2014).

We thus investigated whether Crk plays a role in the collective movements that drive embryonic wound closure, by quantifying wound healing dynamics in control versus *crkS-RNAi* embryos (Figure 10A,B). We used laser ablation to induce wounds in the ventral epidermis of stage 14-15 embryos expressing E-cadherin::GFP, which outlines cells and allows us to visualize cell behavior during wound closure. In wildtype embryos, wounds close rapidly, with almost complete closure after 45 minutes (Fig 10A). Wounds closed more slowly in *crkS-RNAi* embryos (Figure 10B, quantified in C). The rate of wound closure (measured as the decay constant of an exponential fit to the wound area vs. time curve of each wound) decreased by 43% in *crkS-RNAi* embryos with respect to controls (2.5 ± 0.3 hr^1^ versus. 4.4 ± 0.5 hr^1^, respectively) (mean ± SEM). Consistent with this, when we previously measured the rate of closure during the fast phase of wound repair in *abl* maternal/zygotic mutants, it was 37% slower than in wildtype embryos (Zulueta-Coarasa *et al*., 2014). Thus, wound closure provides another example of a morphogenetic process requiring both Abl and Crk.

**Figure 10.**
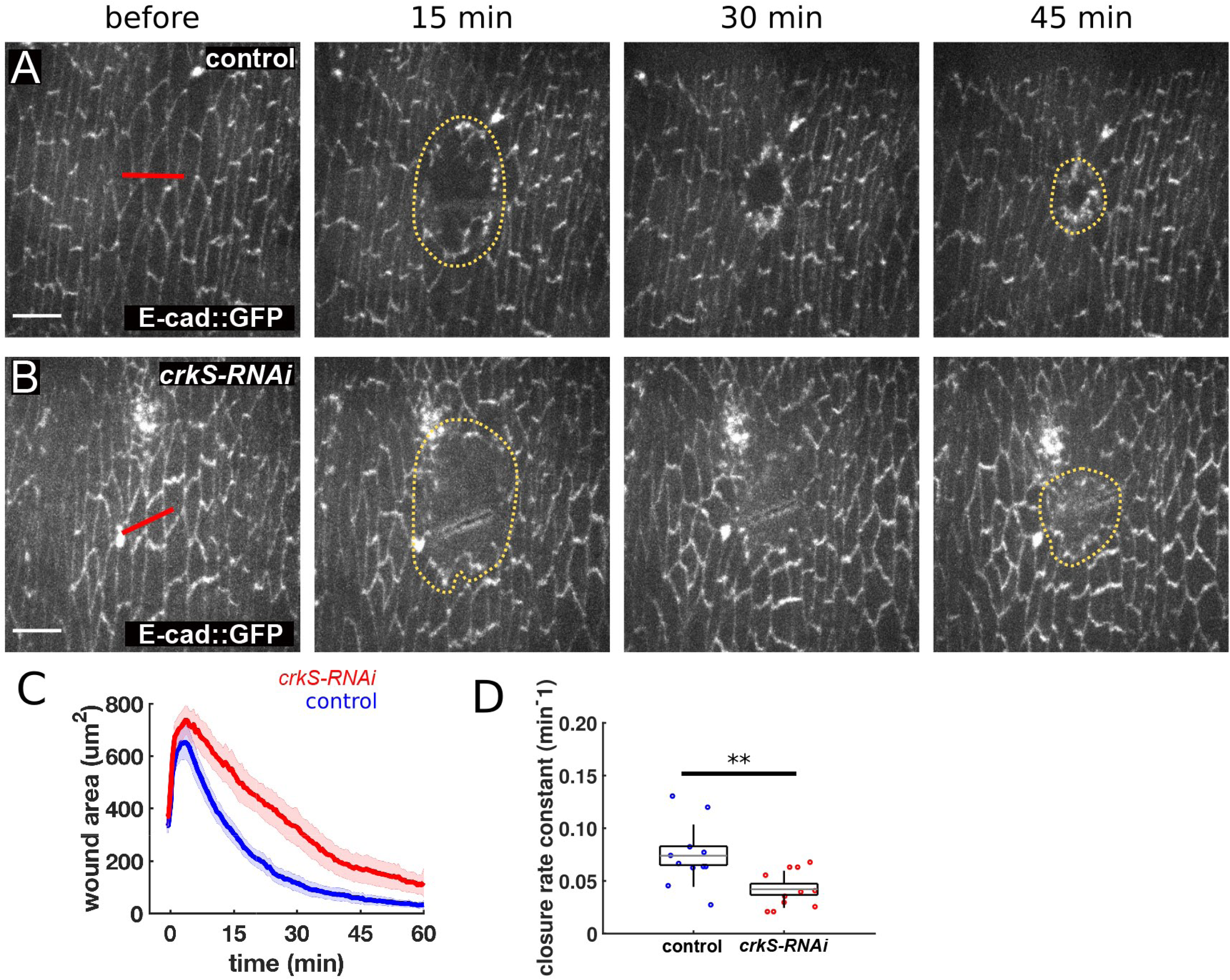
Crk knockdown slows embryonic wound healing. A-B. Wound closure in control (A) and *crkS-RNAi* embryos (B) expressing endo-DE-cadherin::GFP. Red lines indicate wound sites, yellow dotted lines outline the wounds. Anterior, left; ventral, down. Scale bar=10μm. C-D. Mean wound area over time (C) and closure rate constant (D) for controls (blue, n = 11 wounds) and *crkS-RNAi* embryos (red, n = 11). **, P < 0.01 (Mann-Whitney test).

## Discussion

Small adapter proteins play key roles in assembling multiprotein signaling complexes, mediating the interface between cell surface receptors and downstream effectors. The Crk family provides an example. We developed new genetic tools that allowed us to fully assess the role of Crk in embryonic development, revealing novel roles in the cytoskeletal events of syncytial development and cellularization, in central nervous system development, and in embryonic wound healing.

### Crk is an important regulator of actin dynamics during syncytial development and cellularization, acting via SCAR and the Arp2/3 complex

The *Drosophila* syncytial and cellularizing blastoderm provides a superb model for studying dynamic cytoskeletal rearrangements. During each of the last four nuclear cycles, the actin cytoskeleton undergoes a dynamic set of rearrangements. The sequential formation of actin caps and pseudocleavage furrows are critical to allow the underlying nuclei to accurately segregate chromosomes. The subsequent coordinated movement of actin rings around each nucleus and their incomplete closure during cellularization leads to the formation of the initial polarized epithelium. Our data reveal that these coordinated events require Crk function and suggest that defects in actin cap expansion and pseudocleavage furrow invagination after Crk knockdown lead to defects in separation of mitotic spindles and subsequent nuclear loss.

During the normal syncytial division cycles actin caps expand and collide, with collision driving furrow ingression (Zhang *et al*., 2018). Growth of the actin caps requires the DOCK-family guanine nuclear exchange factor Sponge (Postner *et al*., 1992), the Arp2/3 regulator SCAR (Zallen *et al*., 2002), and the Arp2/3 complex (Stevenson *et al*., 2002). Data also support a role for the small GTPase Rac, downstream of Sponge and upstream of SCAR (Zhang *et al*., 2018). The known physical interactions of Crk with DOCK family members and its role in DOCK/ELMO/Rac1 pathways in phagocytic clearance, nervous system remodeling, thoracic closure, and border cell migration are thus intriguing in this regard. The roles for Crk we documented in enhancing cap expansion and furrow invagination are consistent with it playing a role in this pathway, as are the effects of Crk knockdown on levels of SCAR, Arp3, and branched F-actin in actin caps and furrows. Thus, Crk may act upstream of SCAR, although roles in regulating other important players like the formin Diaphanous, Anillin, or the septin Peanut have not been ruled out (Fares *et al*., 1995; Field and Alberts, 1995; Adam *et al*., 2000; Afshar *et al*., 2000; Field *et al*., 2005). Our data are also consistent with a role for Crk in cellularization, though at this stage the exact mechanisms involved and the extent to which this is a primary role or a secondary effect of defects in syncytial divisions remain unclear.

Much work remains to clearly define the pathways and mechanistic connections driving actin rearrangements during syncytial development. For example, the upstream, initiating regulators of this process have yet to be identified. Unlike in myoblast fusion or in RTK-dependent actin remodeling, there is no known role for transmembrane receptors in actin remodeling during syncytial development. Centrosomes play an important role, and can continue to organize actin cap and furrow dynamics even in the absence of nuclei (Raff and Glover, 1989; Stevenson *et al*., 2001), but the mechanisms by which they communicate with the DOCK-family GEF Sponge and ultimately activate SCAR remain unknown. While recent data is consistent with Rac being a downstream effector (Zhang *et al*., 2018), other data suggest Rap1 acts downstream of Sponge during cellularization (Schmidt *et al*., 2018), as it does in later embryonic events (Biersmith *et al*., 2015). Intriguingly, C3G, a Rap1 guanine-nucleotide exchange factor, was the first Crk SH3-domain interacting protein identified (Knudsen *et al*., 1994; Tanaka *et al*., 1994). Src/Crk/C3G/Rap1 signaling is involved in activation of Cdc42 during nectin-induced adherens junction formation (Fukuyama *et al*., 2005). In T-cells, Rap1 activation and integrin-dependent adhesion downstream of T-cell receptor engagement is dependent on both Abl activity and CrkL/C3G recruitment to the plasma membrane (Nolz *et al*., 2008). It will be exciting to explore these mechanistic connections further, and to examine potential roles of other pathway members like Verprolin 1 (Kim *et al*., 2007), and its upstream regulator Blown fuse (Jin *et al*., 2011).

### Crk plays additional roles in the central nervous system and in wound healing

We suspected Crk’s roles would not be restricted to early events, but the major defects in syncytial development and cellularization posed a challenge in determining whether defects in later events were a primary or secondary consequence. For example, we suspect the major disruption of the central nervous system seen in a subset of Crk knockdown embryos reflects early cell loss of both epidermal and neural precursors. This is supported by the strong, though not complete, correlation between defects in the central nervous system (CNS) and defects in the overlying epidermis. In the future it will be important to directly examine stages when neuroblasts invaginate to test this hypothesis. However, our range of knockout and RNAi tools helped circumvent this difficulty. Another subset of embryos, particularly those in the *crkW-RNAi* background, had more mild CNS defects even when the overlying epidermis was intact. These data are consistent with Crk playing a more direct role in axon guidance. Intriguingly, *abl* (Rogers *et al.,* 2016), *SCAR* (Zallen *et al*., 2002), and *sponge/mbc* loss (Biersmith *et al*., 2011) all can lead to CNS defects similar to those seen after Crk knockdown. Crk knockdown also modifies photoreceptor axon targeting defects arising from overexpression of Eyes absent (Hoi *et al*., 2016), an event also regulated by Abl (Xiong *et al*., 2009). Our data also suggest that not all Robo-positive axons respect the normal midline repulsive signals after Crk knockdown, although basic Robo and Slit expression patterns are intact. Future work should directly explore the role of Crk in axon guidance and examine genetic interactions between *crk* and known players in this process. Development of tools to target Crk for proteasomal destruction in specific tissues (e.g., Caussinus and Affolter, 2016)) would be particularly valuable. For example, tissue-specific degradation of Crk could provide new insight into its potential roles in myoblast fusion or axon patterning.

Our examination of embryonic wound healing provided a second place where Crk plays a role post-gastrulation. Wound healing requires a complex interplay between cell adhesion and the actomyosin cytoskeleton (reviewed in Hunter and Fernandez-Gonzalez, 2017). Actin and myosin rapidly accumulate at the wound edge, forming a supracellular contractile cable that coordinates cell movements to help drive rapid wound closure. In parallel, cell-cell junctions are re-modeled, with concentration of the cadherin-catenin complex at tricellular junctions along the wound edge. Our analysis revealed that Crk knockdown slows wound healing. Intriguingly, Abl loss has a similar effect, and in the case of Abl its loss impairs both actin organization at the wound edge and enrichment of the cadherin-catenin complex at tricellular junctions (Zulueta-Coarasa *et al*., 2014). SCAR is also critical for embryonic wound healing (Matsubayashi *et al*., 2015). In the future it will be important to explore which cell biological events required for wound healing are regulated by Crk. Germband retraction is a third post-gastrulation event our data suggest requires Crk. Cell-matrix adhesion plays a key role in this process, in which different tissues exert mechanical force upon one another (reviewed in Lacy and Hutson, 2016). Once again, Abl loss has a parallel effect (Grevengoed *et al*., 2001) to that of loss of Crk. This will provide another place where the cell biological roles of Crk can be explored.

## Methods and Materials

### Fly husbandry and cuticle preparation

All stocks were maintained on standard cornmeal agar media at room temperature or 25°C. All RNAi crosses and controls were maintained at 25°C. Stocks used are described in Table 1. We used RNAi to generate embryos with reduced maternal and zygotic Crk as follows. We used two independent UAS-driven shRNA lines targeting different regions of crk, developed by the Transgenic RNAi Project (TRiP; (Perkins *et al*., 2015))—HMC03964 (use of which is referred to as *crkS-RNAi)* and HMJ22995 (use of which is referred to as *crkW-RNAi)*. Both shRNA constructs are in a VALIUM20 backbone and are inserted in the same genomic locus (attP40) using PhiC31-mediated integration, making them directly comparable. We generated females carrying two copies of *matα*-tubulin-GAL4 (one copy on the second and one on the third chromosome) and a single copy of the indicated UAS-shRNA construct. These females were setup in cups with males either homozygous (*crkS-RNAi*) or heterozygous (*crkW-RNAi*) for the indicated UAS-shRNA and allowed to lay eggs for the appropriate period. For mNG::3XFLAG::Crk overexpression, we generated females carrying two copies of *matα*-tubulin GAL4 and a single copy of UASp-mNG::3XFLAG::dCrk. These females were mated to homozygous UASp-mNG::3XFLAG::dCrk males and allowed to lay eggs for the appropriate period. For simultaneous knockdown/overexpression, we generated a second chromosome bearing both UASp-mNG::3XFLAG::dCrk and UAS-*crkS-RNAi* using standard recombination and generated embryos as described above. To generate *crk* germline clones, we generated females that were heterozygous for the indicated FRT-flanked rescue construct over *crk^ΔattP^* and were simultaneously carrying a single copy of *ovo*-FLP. These females were crossed to *y, w; crk^ΔattP^/In(4) ci^D^ ci^D^ pan^ciD^* males in cups and allowed to lay eggs for the appropriate period. Cuticle preparations were performed as described in Wieschaus and Nüsslein-Volhard (1986)

### Generation of Crk antibody

The ORF from *crk* isoform RA was PCR-amplified from a gBlocks Gene Fragment (IDT, Coralville, IA, USA) using the primers listed below and cloned into the NdeI/EcoRI sites of pET28P (6XHis-dCrk) or the BamHI/EcoRI sites of pGEX-6P-2 (GST-dCrk). Resulting clones were screened by restriction digest and verified using Sanger sequencing.

PCR Primers (restriction sites in bold, target sequence underlined)

pET28P Forward: 5’-ACTG**CATATG**<u>GATACATTTGACGTTTCTGA</u>-3’

Reverse: 5’-ACTG**GAATTC**<u>TTAGCATATTTCTGTGGAGTTTTTG</u>-3’

pGEX-6-P2 Forward: 5’-ACTG**GGATCC**<u>ATGGATACATTTGACGTTTCTGA</u>-3’

Reverse: 5’-ACTG**GAATTC**<u>TTAGCATATTTCTGTGGAGTTTTTG</u>-3’

Recombinant protein was expressed in BL21 Star (DE3) cells and purified using standard affinity chromatography techniques. Pocono Rabbit Farm & Laboratory, Inc. (Canadensis, PA, USA) raised antisera against 6XHIS-dCrk in two rabbits. Antisera were affinity-purified against GST-dCrk using standard techniques prior to use for Western blot analysis.

### Generation of a *crk* null allele

The *crk* null allele, *crk^ΔattP^*, was generated by using CRISPR to replace the entire *crk* locus (beginning 110bp upstream of transcription start site and ending 99bp beyond the 3’UTR) with a 50 base-pair attP phage recombination site and positive marker (a loxP-flanked 3xP3-DsRed cassette) as described in (Gratz *et al*., 2014). Briefly, guide RNAs flanking the *crk* locus were selected using flyCRISPR Target Finder (http://tools.flycrispr.molbio.wisc.edu/targetFinder/) to identify 19-20bp CRISPR targets with NGG PAM sites and zero predicted off-target effects using maximum stringency. Annealed gRNA oligo nucleotides were cloned into the BbsI site of pU6-BbsI-chiRNA. We generated a dsDNA donor template for homology-directed repair by cloning upstream and downstream flanking regions of the crk locus into the AarI and SapI sites of pHD-DsRed-attP, respectively. An 815bp upstream homology region, extending from 5’-CTTGAGCATGCAAAGGAATG-… to …GCTGACAGTACGTCCT-3’ and flanked by AarI sites was synthesized as a gBlocks Gene Fragment (IDT, Coralville, IA, USA). A 1kb downstream homology region was PCR amplified from genomic DNA using the primers described below. An injection mixture containing both U6-gRNA plasmids (75ng/ul each) and the dsDNA template (250ng/ul) was injected into *y^1^, M{vas-Cas9.RFP-}ZH-2A, w^1118^*flies. Injections performed by BestGene, Inc. (Chino Hills, CA, USA). Surviving G_0_ animals were outcrossed to *y w* and CRISPR edited lines were identified by the presence of DsRed eye fluorescence. DsRed positive G_1_ animals were further outcrossed to *y w* to remove *M{vas-Cas9.RFP-}ZH-2A* and subsequently established as stocks over *In(4) ci^D^, ci^D^ pan^ciD^*. Deletion of the *crk* locus was verified by PCR using the primers described below.

<u>Oligo nucleotides for gRNA1:</u>

Target (PAM underlined): GCTGACAGTACGTCCTAAA<u>GGG</u>

Sense oligo: 5’-CTTCGCTGACAGTACGTCCTAAA-3’

Antisense oligo: 5’-AAACTTTAGGACGTACTGTCAGC -3’

<u>Oligo nucleotides for gRNA2:</u>

Target (PAM underlined): GAATTGTGTAACTAAACGGT<u>AGG</u>

Sense oligo: 5’-CTTCGAATTGTGTAACTAAACGGT-3’

Antisense oligo: 5’-AAACACCGTTTAGTTACACAATTC-3’

<u>Primers for PCR amplification of downstream homology region (Sap1 site in bold, target sequence underlined):</u>

Forward: 5’-GACT**GCTCTTC**ATAT<u>TGCTACAGAAAACCCTTTGG</u> -3’

Reverse: 5’- GACT**GCTCTT**CAGAC<u>GAACATAGACAGCACAAGTTTGTTG</u> -3’

<u>Primers for PCR verification of *crk* locus deletion:</u>

Detection of DsRed:

Forward: 5’-CGAGGACGTCATCAAGGAGT-3’

Reverse: 5’-GGTGATGTCCAGCTTGGAGT-3’

Flanking upstream HR:

Forward: 5’-AATTGCGCGCTGTTAAAAAT-3’

Reverse: 5’-GCTTCGAGCCGATTGTTTAG-3’

Flanking Downstream HR:

Forward: 5’-GTGGACTCCAAGCTGGACAT-3’

Reverse: 5’-TAACAGGCAAAAACGCTTCC-3’

### Generation of rescue constructs for PhiC31 integrase-mediated targeting into *crk^ΔattP^*

The following rescue constructs were generated by introducing the indicated PCR amplified fragments into EcoRI/KpnI linearized pGE-attB^GMR^ ((Huang *et al*., 2009) using NEBuilder HiFi DNA Assembly (New England Biolabs). We generated a genomic rescue construct by PCR amplifying a 3,151bp fragment of the *crk* locus, extending from 5’-AAAGGGGTTTACCACCACAG-3’… to …5’- TTGTGTAACTAAACGGTAGG-3’ from genomic DNA—this rescue construct corresponds to the exact sequence deleted in generation of *crk^ΔattP^*.

<u>Primers used (overlap with pGE-attB^GMR^ in lowercase, target sequence underlined):</u>

Forward: 5’-gctccccgggcgcgtactccacgaattc<u>AAAGGGGTTTACCACCAC</u>-3’

Reverse: 5’-attatacgaagttatggtac<u>CCTACCGTTTAGTTACACAATTC</u>-3’

We generated a cDNA-based rescue construct using the ORF from *crk* isoform RA by assembling the following PCR amplified fragments into EcoRI/KpnI linearized pGE-attB^GMR^:

5’ deleted region (includes 5’UTR and 110bp upstream of the transcription start site):

Primers used (overlap with neighboring fragment in lowercase, target sequence underlined):

Forward: 5’- gggctccccgggcgcgtactccacgaattc<u>AAAGGGGTTTACCACCAC</u>-3’

Reverse: 5’- tgtcttcttcacctttggaaaccat<u>TTCCGTTTAAGGTTTATCTTAGAC</u>-3’

mNG::3XFLAG::dCrk (synthesized as a gBlocks Gene Fragment (IDT, Coralville, IA, USA), 3XFLAG sequence in bold, dCrk-RA sequence underlined):

ATGGTGAGCAAGGGCGAGGAGGATAACATGGCCTCTCTCCCAGCGACACATGAGTTACACAT CTTTGGCTCCATCAACGGTGTGGACTTTGACATGGTGGGTCAGGGCACCGGCAATCCAAATGATG GTTATGAGGAGTTAAACCTGAAGTCCACCAAGGGTGACCTCCAGTTCTCCCCCTGGATTCTGGTC CCTCATATCGGGTATGGCTTCCATCAGTACCTGCCCTACCCTGACGGGATGTCGCCTTTCCAGGC CGCCATGGTAGATGGCTCCGGCTACCAAGTCCATCGCACAATGCAGTTTGAAGATGGTGCCTCCC TTACTGTTAACTACCGCTACACCTACGAGGGAAGCCACATCAAAGGAGAGGCCCAGGTGAAGGGG ACTGGTTTCCCTGCTGACGGTCCTGTGATGACCAACTCGCTGACCGCTGCGGACTGGTGCAGGTC GAAGAAGACTTACCCCAACGACAAAACCATCATCAGTACCTTTAAGTGGAGTTACACCACTGGAAA TGGCAAGCGCTACCGGAGCACTGCGCGGACCACCTACACCTTTGCCAAGCCAATGGCGGCTAAC TATCTGAAGAACCAGCCGATGTACGTGTTCCGTAAGACGGAGCTCAAGCACTCCAAGACCGAGCT CAACTTCAAGGAGTGGCAAAAGGCCTTTACCGATGTGATGGGCATGGACGAGCTGTACAAG**ATGG ACTACAAAGACCATGACGGTGATTATAAAGATCATGACATCGATTACAAGGATGACGATGACAA G**<u>ATGGATACATTTGACGTTTCTGATAGGAACAGCTGGTACTTTGGTCCCATGTCTAGACAGGATGC</u> <u>TACTGAAGTTTTGATGAACGAACGCGAGCGGGGAGTGTTTTTAGTCCGTGATAGTAACTCGATAGC AGGGGATTATGTACTTTGTGTAAGAGAAGATACAAAAGTTAGCAACTACATCATTAACAAAGTTCAA</u> <u>CAACAGGATCAAATCGTTTACCGCATTGGGGATCAGTCTTTTGACAATCTACCGAAACTCTTAACTT</u> <u>TTTACACTCTTCATTATTTGGATACAACCCCTTTAAAACGGCCTGCGTGTAGAAGGGTGGAAAAAG</u> <u>TAATAGGAAAGTTCGATTTCGTTGGCAGCGATCAAGATGATTTACCTTTTCAAAGAGGTGAAGTTTT</u> <u>AACAATAGTTCGAAAAGACGAGGATCAATGGTGGACTGCGCGTAACTCCTCGGGGAAAATTGGTC</u> <u>AAATACCGGTTCCCTATATACAACAGTATGACGATTATATGGATGAAGATGCTATTGATAAAAACGA</u> <u>ACCTTCCATTTCGGGATCTAGCAATGTATTTGAAAGTACTCTTAAAAGGACAGATTTAAATCGAAAA</u> <u>CTACCTGCATACGCCCGCGTAAAACAGTCAAGGGTCCCTAACGCATACGATAAGACTGCATTAAAA</u> <u>TTGGAAATAGGTGACATTATTAAAGTCACTAAAACAAACATTAATGGGCAATGGGAGGGAGAATTA</u> <u>AATGGAAAAAATGGTCATTTTCCCTTCACGCACGTTGAATTTGTCGATGATTGTGATTTAAGCAAAA</u> <u>ACTCCACAGAAATATGCTAA</u>

<u>3’ deleted region (includes 3’UTR and 99bp downstream):</u>

Primers used (overlap with neighboring fragment in lowercase, target sequence underlined):

Forward: 5’- caaaaactccacagaaatatgctaa<u>ATAGGAAGGAATGGAAGG</u>-3’

Reverse: 5’- catacattatacgaagttatggtac<u>CCTACCGTTTAGTTACACAATTC</u>-3’

For our FRT-flanked rescue constructs, we introduced flanking FRT sites 104bp upstream of the transcription start site and 99bp downstream of the end of the 3’UTR using a Q5 Site-Directed Mutagenesis kit (New England Biolabs) and the following primers:

<u>Upstream FRT (FRT sequence in lowercase, target sequence underlined):</u>

Forward: 5’-tagagaataggaacttc<u>GTTTACCACCACAGAGCAG</u>-3’

Reverse: 5’-gaaagtataggaacttca<u>CCCTTTGAATTCGTGGAG</u>-3’

<u>Downstream FRT (FRT sequence in lowercase, target sequence underlined):</u>

Forward: 5’-tagagaataggaacttc<u>GTACCATAACTTCGTATAATG</u>-3’

Reverse: 5’-gaaagtataggaa<u>CTTCACCTACCGTTTAGTTACAC</u>-3’

All rescue constructs were verified by a combination of PCR, restriction digest, and Sanger sequencing prior to injection. Transgenic lines were generated by co-injecting rescue constructs and PhiC31 helper plasmid into y, w; *crk^ΔattP^*/ *In(4) ci^D^, ci^D^ pan^ciD^* embryos. Injections performed by BestGene, Inc. (Chino Hills, CA, USA).

### Generation of Crk overexpression constructs

We generated two constructs for overexpressing fluorescent protein-tagged Crk using the UAS/GAL4 system—mNG-3XFLAG-Crk and tdTomato-3XFLAG-Crk. mNG-3XFLAG-Crk was generated by cloning PCR amplified mNG-3XFLAG-dCrk into the KpnI/SphI sites of pTIGER (Ferguson *et al*., 2012; Scott Ferguson, Fredonia University). Similarly, tdTomato-3XFLAG-Crk was generated by sequentially cloning PCR amplified tdTomato and 3XFLAG-Crk sequences into the KpnI/NotI and NotI/SphI sites of pTIGER, respectively. Constructs were verified using both restriction digest and Sanger sequencing. Transgenic lines were generated using PhiC31 integrase-mediated site-specific integration into attP16 on 2R (Bloomington Stock #53C4; *y*^1^, *w*^67c23^; P{CaryP}attP16). Injections were performed by BestGene, Inc. (Chino Hills, CA, USA).

<u>Primers used:</u>

mNG::3XFLAG::Crk (restriction sites in bold, Kozac consensus sequence in lowercase, target sequence underlined)

Forward: 5’-GACT**GGTACC**cacc<u>ATGGTTTCCAAAGGTGAAGAA</u>-3’

Reverse: 5’-ACTG**GCATGC**<u>TTAGCATATTTCTGTGGAGTTTTTG</u>-3’

3XFLAG::Crk (restriction sites in bold, target sequence underlined)

Forward: 5’- GACT**GCGGCCGC**A<u>ATGGACTACAAGGATCATGACGG</u>-3’

Reverse: 5’-ACTG**GCATGC**<u>TTAGCATATTTCTGTGGAGTTTTTG</u>-3’

tdTomato (restriction sites in bold, Kozac consensus sequence in lowercase, target sequence underlined)

Forward: 5’-GACT**GGTACC**cacc<u>ATGGTGAGCAAGGGCGAG</u> -3’

Reverse: 5’-ACTG**GCGGCCGC**<u>CTTGTACAGCTCGTCCATGC</u> -3’

### Western blotting

Flies were allowed to lay eggs on apple juice agar plates with yeast paste for the indicated time. Dechorionated embryos were lysed by manual homogenization in ice-cold RIPA lysis buffer [1% NP-40; 0.5% Na deoxycholate; 0.1% SDS; 50mM Tris, pH 8; 300mM NaCl; 1X Halt protease and phosphatase inhibitor cocktail (Thermo Scientific)]. Lysates were cleared by centrifugation at 21,000g at 4°C for 1 min and protein concentration determined using Bio-Rad Protein Assay Dye. Normalized samples were mixed 1:1 with 2X SDS Sample Buffer and boiled for 5 min. Samples were either run immediately or stored at −20°C and boiled 5 min prior to running on 10% SDS-PAGE and transferring to nitrocellulose. Blots were blocked in 5% milk in TBST (0.1% Tween20 in Tris-buffered saline, pH 7.6). Western blots were probed with mouse anti-*γ*Tubulin (1:2,500) (GTU-88, Sigma Aldrich), and Rabbit anti-dCrk (1:10,000). Secondary antibodies used were IRDye-conjugated donkey anti-mouse (1:5,000) and goat anti-rabbit (1:5,000). Blots were imaged using Odyssey CLx infrared imaging system (LI-COR Biosciences). Band densitometry performed using ImageStudio software version 4.0.21 (LI-COR Biosciences). Statistical comparisons were made using unpaired t-tests in Prim8 (GraphPad).

### Immunofluorescence and image analysis

Flies were allowed to lay eggs for the appropriate time for the desired stages of development on apple juice/agar plates with yeast paste. Embryos were dechorionated in 50% bleach, washed 3X in 0.1% Triton-X, and fixed at room temperature in 1:1 heptane/formaldehyde diluted in phosphate-buffered saline (PBS). For phalloidin staining, syncytial embryos were fixed for 30min using 18.5% formaldehyde and devitellinized by either hand peeling or shaking in 1:1 heptane/90% ethanol. In all other instances, embryos were fixed for 20min using 4% formaldehyde and devitellinized by shaking in 1:1 heptane/methanol. Embryos were blocked in PNT (PBS, 0.1% Triton-X100, and 1% normal goat serum) for ≥30 min followed by incubation with primary antibodies diluted in PNT at room temperature for 4 h or overnight at 4°C. Following washing three times with PBT (PBS, 0.1% Triton-X100), embryos were incubated in secondary antibodies diluted in PNT at room temperature for 2 h or overnight at 4°C. Embryos were washed three times with PBT and mounted on glass slides using Aquapolymount (Polysciences, Warrington, PA). Imaging was done on a Zeiss (Oberkochen, Germany) LSM-5 Pascal, a Zeiss 710, or a Zeiss 880 scanning confocal microscope. Primary antibodies used were: anti-BP102 (1:200), anti-FasII (1:100), anti-Robo (1:100), anti-SCAR (1:50), anti-Slit (1:10; all from the Developmental Studies Hybridoma Bank, Iowa City, IA); anti-Arp3 (1:500; (Stevenson *et al*., 2002)), anti-pTyr (1:1,000; 4G10 clone; EMD Millipore); and anti-*α*Tubulin (1:2,000; DM1A clone; Sigma). Secondary antibodies used were: Alexa Fluor 488, 568, and 647 conjugated Goat anti-mouse (pan or isotype-specific as required), Goat anti-rat, or Goat anti-rabbit secondaries (1:500; Life Technologies). Other stains used: Alexa Fluor 488, 568, or 647 conjugated phalloidin (1:500; Life Technologies), DR (1:500; BD Biosciences), and AF647 wheat germ agglutinin (1:1,000; Life Technologies).

### Analysis of SCAR and Arp3 localization

To be able to directly compare fluorescent intensity levels, wild-type control (Histone::RFP) and crkS-RNAi embryos were collected and processed in the same tube. Embryo fixation and staining carried out as described above in Immunofluoresence and image analysis. Embryos were fixed in 18.5% formaldehyde diluted in PBS for 30min at room temperature and devitellinized using 1:1 heptane/90% ethanol. Images were acquired as 8-bit, single plane confocal sections using a Zeiss (Oberkochen, Germany) LSM-5 Pascal. All images within a given experiment were acquired using the same settings (i.e., laser power, gain, offset, etc.). These confocal sections were taken at the level in which actin caps or pseudocleavage furrows appeared most obvious for each embryo. To measure fluorescent intensity in caps, we used the Fiji distribution of ImageJ (version 1.52N; (Schindelin *et al*., 2012; Rueden *et al*., 2017)) to draw 30 pixel diameter circles, centered over 10 randomly selected caps for each embryo analyzed and recorded mean gray values at each position. To measure fluorescent intensity in pseudocleavage furrows, we freehand drew 3-pixel wide lines over 10 randomly selected pseudocleavage furrows and recorded mean gray values at each position. We compared fluorescent intensity levels for SCAR, Arp3, and phalloidin between genotypes by plotting the mean gray values for each individual cap or pseudocleavage furrow measured. Statistical comparisons made using unpaired t-tests in Prism8 (GraphPad).

### Wound healing assays

Crk knockdown was achieved by crossing *mat67*, *endo-D-E-cadherin:GFP*; *mat15* virgins to *crkS-RNAi*/*CyO*; *Pri*/*TM3* males. *mat67*, *endo-DE-cadherin:GFP*/*crkS-RNAi*; *mat15*/(*Pri* or *TM3*) virgins were subsequently crossed to *crk*S*-RNAi* males, and their progeny were wounded and imaged. The progeny of *mat67*, *endo-DE-cadherin:GFP*/*CyO*; *mat15*/(*Pri* or *TM3*) virgins crossed to *y, w* males were used as controls. For wound healing assays, stage 14-15 embryos were dechorionated for two minutes in 50% bleach. Embryos were arranged on an apple juice-agar pad with the ventral side down and transferred to a coverslip coated with heptane glue. Embryos were then covered with a 1:1 mix of halocarbon oil 27:700 (Sigma Aldrich). Embryos were imaged using a Revolution XD spinning disk confocal microscope (Andor Technology) with an iXon Ultra 897 camera (Andor Technology), a 60X (NA 1.35; Olympus) oil-immersion lens, and Metamorph software (Molecular Devices). Wounding was performed using a Micropoint nitrogen dye cell laser (Andor Technology) tuned to 365 nm. To wound the embryos, 10 laser pulses were delivered at spots 2 μm apart along a 14 μm line. 16-bit Z-stacks consisting of 16 slices at 0.2 μm steps were acquired every 30 seconds after wounding. Maximum intensity projections of time-lapse images were analyzed using SIESTA (Fernandez-Gonzalez and Zallen, 2011), custom software written in MATLAB (Mathworks) using the DIPImage toolbox (Delft University of Technology). The wound margin was segmented using the semi-automated LiveWire tool at every time point within the time-lapse image. A two-parameter exponential function was fitted to individual wound healing curves:

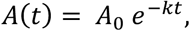

where *A*(*t*) represents the area of the wound at time *t*, *A*_0_ is the maximum wound area, and *k* is the decay constant of the exponential. A least-squares fit by the Gauss-Newton method was done starting at the point of maximum wound area. The decay constant of the exponential was used to quantify the rate of wound closure.

### Live imaging

Embryos with Crk knockdown and carrying live imaging constructs were produced by crossing either *crkS-RNAi / CyO; His2av mRFP, sGMCA* or *crkS-RNAi / CyO; MRLC::GFP* males to *matα tub* - GAL4 *II; matα tub* -GAL4 *III* females. *crkS-RNAi/ matα tub* -GAL4 II*; His2av mRFP, sGMCA/matα tub* GAL4 III or *crkS-RNAi/ matα tub* -GAL4 II*; MRLC::GFP/matα tub* GAL4 III female progeny were then crossed to *crkS-RNAi* males and the progeny were imaged. *CyO/matα tub* -GAL4 II*; His2av mRFP, sGMCA/matα tub*-GAL4 III females crossed to *y w* males were used as controls. Flies were allowed to lay on apple juice agar plates with yeast for 2 hours. Embryos were dechorionated in 50% bleach for 3 minutes, washed 3X with .01% Triton-X and transferred to a fresh apple juice/agar plate where they were immersed in halocarbon oil 27(Apex Bioresearch Products). 7-10 embryos were then transferred to halocarbon oil on the gas permeable membrane of a Lumox dish (Sarstedt) and covered with a glass coverslip. Imaging was done on a Zeiss LSM-5 Pascal scanning confocal microscope (Oberkochen, Germany). Embryos were imaged at the apical cortex where the tops of actin caps were visible. Frames were taken at 10 second intervals for about 2 hours or until cellularization began.

### Furrow Analysis

The length of 5 metaphase furrows per embryo were measured using the Fiji distribution of ImageJ (version 1.52N; (Schindelin *et al*., 2012; Rueden *et al*., 2017)). Furrows were selected by drawing a perpendicular line between two unique spindles that had not collided with another, such that the line passed through the middle of the dividing furrow. The ImageJ Stacks>Reslice function was used to construct a Z-axis projection along that line. The resulting projected image was binarized using the Max Entropy Auto Threshold feature (Kapur *et al*., 1985) to improve visibility of the furrow. Furrows were then measured by drawing a line from the top of the furrow to the bottom. In some cases when not enough of a furrow could be discerned to be effectively measured but some pixels were still visible, a value of 0.54μm was recorded as this is the step length between slices of the z stack.

### Cap Expansion Analysis

Cap expansion during nuclear cycle 10 was quantified in 3 non-adjacent caps per embryo live imaged as above. ImageJ was used for the analysis. Videos were binarized using the Default Auto Threshold feature and processed to remove some noise using Process>Noise> Despeckle to improve visibility of cap borders. Cap area was measured by freehand outlining the cap at each frame, from when they first appear at the cortex to when they began to break up to enter the next nuclear cycle. Expansion time was reported as the time in seconds between when the cap first was measurable to when it was maximally expanded. Maximum expansion was defined as the maximum area of the cap before expansion plateaued or decreased. Expansion rate was calculated as the difference in area between when the cap was first measured to when it was maximally expanded divided by the expansion time. Cycle length was measured as the seconds between when caps first reappear to when they appear in the next cycle

## Acknowledgements

We are grateful to Victoria Deneke and Stefano Di Talia for advice and guidance in live imaging, Yixie Zhang, Tony Harris, Steve Rogers and Kevin Slep for advice and helpful discussions, to the Bloomington *Drosophila* Stock Center, the Transgenic RNAi Project (TRiP) and the Developmental Studies Hybridoma Bank for reagents, to Andrew Hudson and Lynn Cooley for the Arp3 antisera, to Scott Ferguson for pTIGER vectors, to Tony Perdue of the Biology Imaging Center, and to members of the Peifer lab for helpful discussions and reading the manuscript. The work was supported by NIH R35 GM118096 to M.P. A.S. was supported by T32 CA009156 and F32 GM117803. A.B. was supported by the UNC Science and Math Achievement and Resourcefulness Track (SMART), in partnership with the North Carolina Louis Stokes Alliance for Minority Participation (NC-SLSAMP; supported by NSF HRD-1202467), and by the William W. and Ida W. Taylor fellowship through the UNC Summer Undergraduate Research Fellowship (SURF) program. R.F.G. and A. F. are supported by grants from the Canada Foundation for Innovation (#30279) and the Canadian Institutes for Health Research (#156279). RFG is the Tier II Canada Research Chair in Quantitative Cell Biology and Morphogenesis.

---

Supplemental Movie 1. Crk knockdown often leads to wide-spread nuclear loss but embryos can largely repair the epithelium before gastrulation. Time-lapse movies showing actin dynamics in control and *crkS-RNAi* embryos expressing Moesin::GFP to label F-actin. The Movie begins during the last syncytial nuclear division and extends until the onset of gastrulation. In the *crkS-RNAi* embryo, large patches of nuclear loss are apparent following the last nuclear division. During cellularization, cells at the margins of these patches can be seen exhibiting protrusive cell behavior and migrating into areas of cell loss to enclose the blastoderm embryo prior to the onset of gastrulation. Images acquired at 10 s intervals and displayed at 10 frames per second. Time between frames varies between genotypes due to different sizes of region scanned. Scan time was 2.82 seconds for the control embryo and 2.30 seconds for the *crkS-RNAi* embryo.

Supplemental Movie 2. Crk knockdown leads to reduced actin cap expansion and less apparent pseudocleavage furrows during syncytial nuclear cycles. Time-lapse movies showing actin dynamics in control and *crkS-RNAi* embryos expressing Moesin::GFP to label F-actin during nuclear cycles 10-14. In both control and *crkS-RNAi* embryos, actin caps form, expand, and merge into pseudocleavage furrows. However, in *crkS-RNAi* conditions, actin cap expansion is reduced compared to controls, leading to impaired pseudocleavage furrow formation. Images acquired every 10 s and displayed at 10 frames per second. Time between frames varies between genotypes due to different sizes of region scanned. Scan time was 2.30 seconds for the control embryo and 2.93 seconds for *crkS-RNAi* embryo. Focal plane was occasionally adjusted to maintain a clear view of the furrows. This video is related to figure 6K,L.

Supplemental Movie 3. Crk loss impairs actin cap expansion and myosin remodeling during syncytial nuclear cycles. Time-lapse movies showing myosin dynamics in control and *crkS-RNAi* embryos during nuclear cycles 10-14, via MRLC::GFP (=Sqh). In control embryos, inter-cap myosin zones are remodeled during actin cap expansion, resulting in an enrichment of myosin in pseudocleavage furrows. In *crkS-RNAi* embryos, this remodeling is impaired, resulting in broader inter-cap myosin zones and reduced enrichment of myosin in pseudocleavage furrows. Images acquired every 10 s and displayed at 10 frames per second. Time between frames varies between genotypes due to different sizes of region scanned. Scan time was 2.01 seconds for the control embryo and 2.43 seconds for the *crkS-RNAi* embryo. Focal plane was occasionally adjusted to maintain a clear view of the furrows. This video is related to figure 6Q-T.

