## Supplemental Figures for "The Crk adapter protein is essential for *Drosophila* embryogenesis, where it regulates multiple actin-dependent morphogenic events"

Supplemental Figure 1. Schematic representation of *crk*$^{\Delta attP}$ and *crk*$^{\Delta attP}$-targeted rescue constructs, including PCR genotyping results used to verify generation of *crk*$^{\Delta attP}$. A. Schematic representation of the wildtype *crk* locus before and after CRISPR/Cas9-mediated deletion of *crk* to generate *crk*$^{\Delta attP}$, with PCR primers used for genotyping indicated (see Methods for details). B. Representative gel image of PCR genotyping results for wildtype and *crk*$^{\Delta attP}$ heterozygotes. A-B. A series of PCR primers were designed such that they would only generate a PCR product if the attP-loxP-3xP3-DsRed cassette was integrated into the *crk* locus at the correct location and orientation (location of primers in A). PCR amplification of genomic DNA extracted from wildtype or *crk*$^{\Delta attP}$ heterozygotes resulted in detectable bands of the expected size only in *crk*$^{\Delta attP}$ heterozygotes, indicating the desired edit was made. Note: lanes in (B) are taken from the same gel and were cut to remove extraneous lanes. C. Schematic representation of *crk* locus following targeted integration of cDNA and genomic rescue constructs into *crk*$^{\Delta attP}$ via PhiC31-mediated recombination are depicted in these schematics (see Methods for details). The plasmid-derived sequences (e.g., *white* selectable marker, Ampicillin resistance, P-element ends), as well as the 3xP3-DsRed cassette could be removed by Cre-mediated recombination, but we have not done so.

Suppl. Figure 1

A

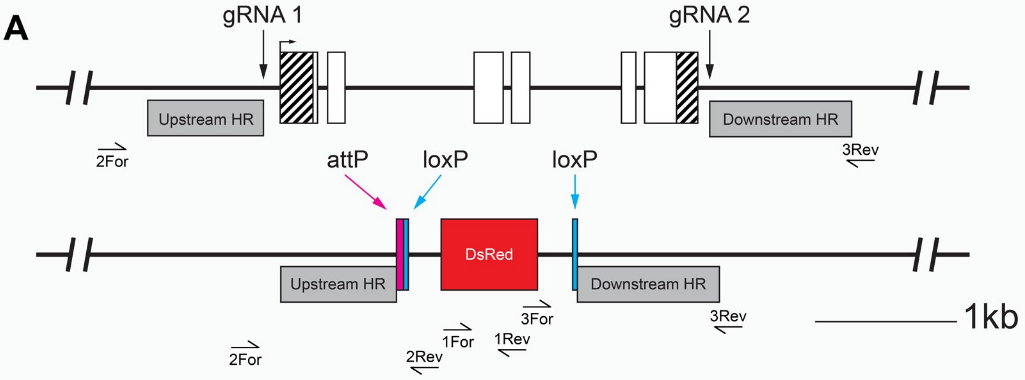


B

Expected product sizes

| Primer | Edited Locus | Wild-type Locus |
|---|---|---|
| 1For<br>1Rev | 595 bp | n/a |
| 2For<br>2Rev | 1,482 bp | n/a |
| 3For<br>3Rev | 1,533 bp | n/a |

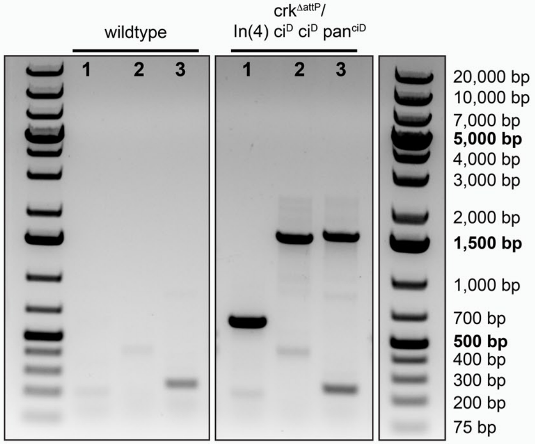


C

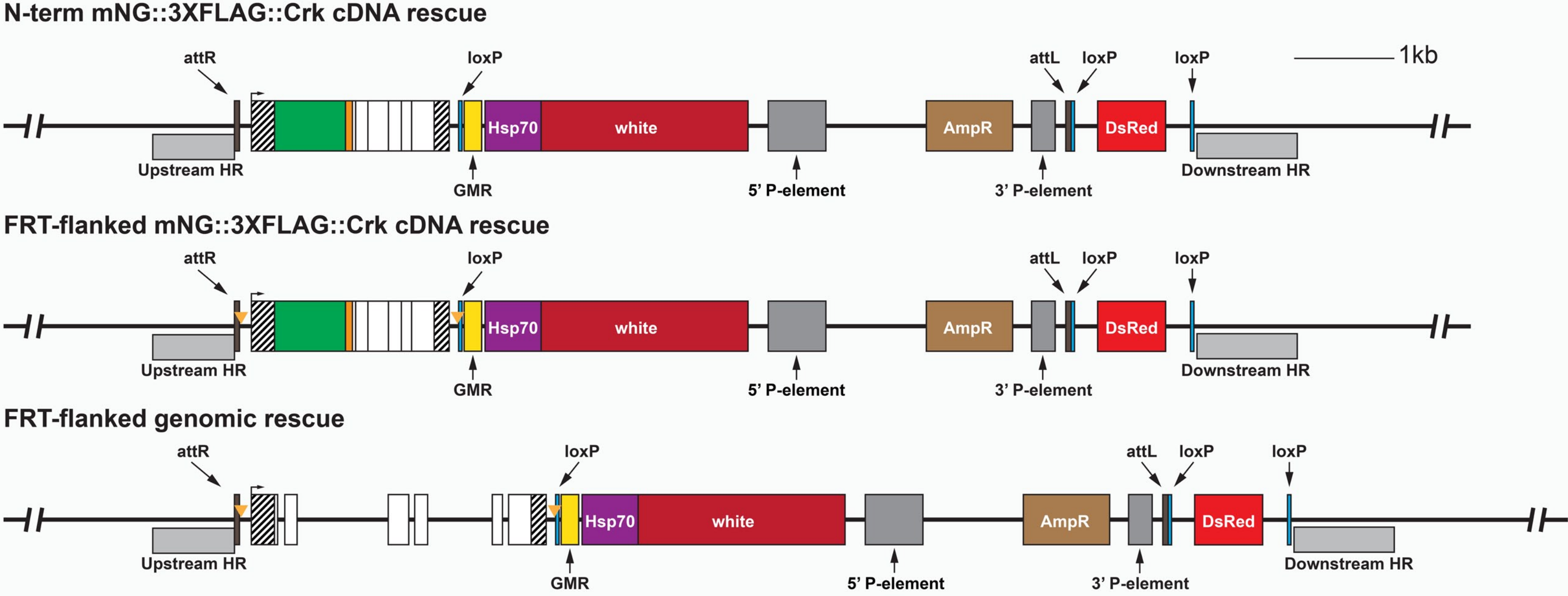

Supplemental Figure 2. Representative examples depicting how fluorescence intensity measurements were made to compare SCAR and Arp3 levels in *crkS-RNAi* embryos vs controls. A-A''. Representative example, actin caps in a wildtype control embryo (Histone::RFP) stained with phalloidin and an antibody against Arp3. B. Quantification of fluorescence intensity of phalloidin and Arp3 in actin caps from embryo in A-A''. C-C''. Representative example of pseudocleavage furrows in a wildtype control embryo (Histone::RFP) stained with phalloidin and an antibody against Arp3. D. Quantification of fluorescence intensity of phalloidin and Arp3 in pseudocleavage furrows from embryo in C-C''. To measure fluorescence intensity of phalloidin and Arp3 in actin caps, we drew 30-pixel diameter circles centered over ten randomly selected caps (A-A'', white circles=regions measured, magenta arrows highlight regions measured) and recorded mean gray values from both channels (B, black bar=mean, error bars=standard deviation). To measure fluorescence intensity of phalloidin and Arp3 in pseudocleavage furrows, we freehand drew 3-pixel wide lines centered over ten random pseudocleavage furrows (C-C'', white lines=regions measured, magenta arrows highlight regions measured) and recorded mean gray values from both channels (D, black bar=mean, error bars=standard deviation). Mean gray values for phalloidin and Arp3 are displayed as scatter plots in B (actin caps) and D (pseudocleavage furrows). The same approach was used to measure fluorescence intensity of F-actin and SCAR levels in actin caps and pseudo cleavage furrows. All scale bars=50µm.

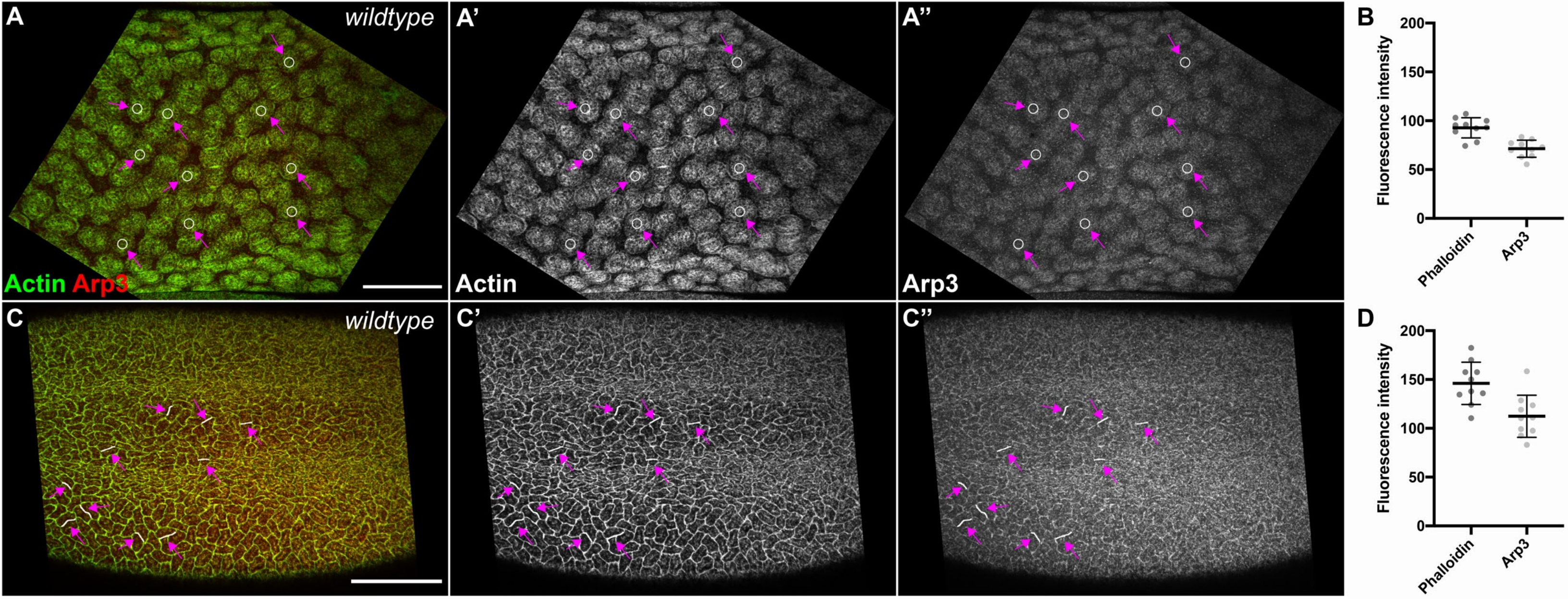
A
wildtype
Actin Arp3
A'
Actin
A''
Arp3
B
Fluorescence intensity
200
150
100
50
0
Phalloidin
Arp3
C
wildtype
C'
C''
D
Fluorescence intensity
200
150
100
50
0
Phalloidin
Arp3

Supplemental Figure 3. Severe defects in CNS patterning in Crk knockdown embryos coincide with disruption of the overlying epidermis. A-E'. Stage 15 embryos, antigens and genotypes indicated. Paired images at the level of the epidermis (left) and the CNS (right). A-A'. Representative example, *crk* knockdown embryo with severe disruption of CNS patterning and strong disruption of epidermal patterning, including multiple large epidermal holes (arrows). B-B'. Representative example, *crk* knockdown embryo in which disruptions of axon patterning lie under regions where there are holes in the epidermis (arrows). C-C'. Representative example, *crk* knockdown embryo with more subtle epidermal defects, such as a denticle belt deletion, with an underlying CNS defect (arrow). D-E". Representative images, *crk* knockdown embryos in which we observed milder CNS defects but the overlying epidermis appears normal (arrows), suggesting at least some of the CNS defects observed could be due to CNS-specific roles for Crk. All scale bars=50μm.

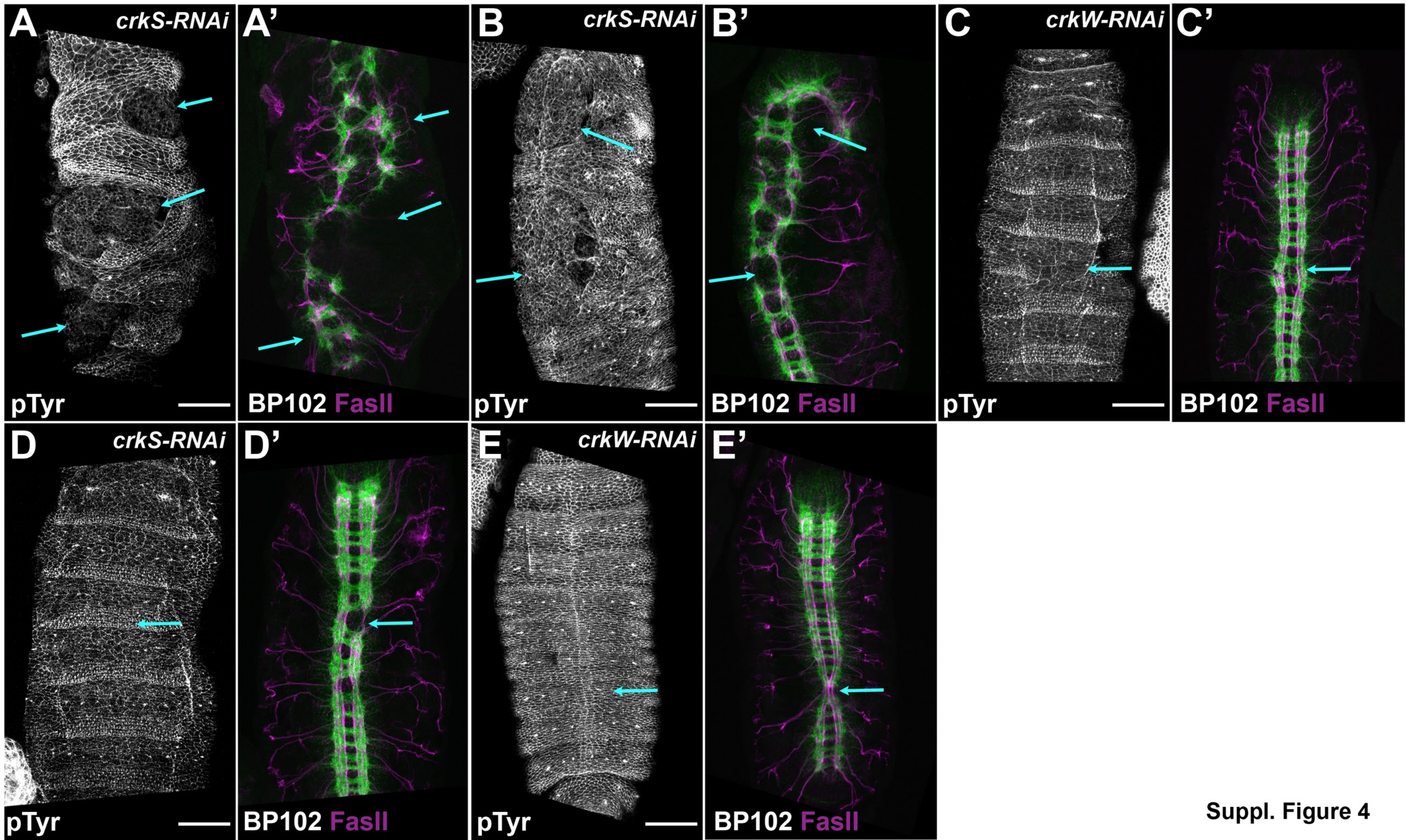


Suppl. Figure 4

**Supplemental Movie Legends**

Supplemental Movie 1. Crk knockdown often leads to wide-spread nuclear loss but embryos can largely repair the epithelium before gastrulation. Time-lapse movies showing actin dynamics in control and *crkS-RNAi* embryos expressing Moesin-GFP to label F-actin. The Movie begins during the last syncytial nuclear division and extends until the onset of gastrulation. In the *crkS-RNAi* embryo, large patches of nuclear loss are apparent following the last nuclear division. During cellularization, cells at the margins of these patches can be seen exhibiting protrusive cell behavior and migrating into areas of cell loss to enclose the blastoderm embryo prior to the onset of gastrulation. Images acquired at 10 s intervals and displayed at 10 frames per second. Time between frames varies between genotypes due to different sizes of region scanned. Scan time was 2.82 seconds for the control embryo and 2.30 seconds for the *crkS-RNAi* embryo.

Supplemental Movie 2. Crk knockdown leads to reduced actin cap expansion and less apparent pseudocleavage furrows during syncytial nuclear cycles. Time-lapse movies showing actin dynamics in control and *crkS-RNAi* embryos expressing Moesin-GFP to label F-actin during nuclear cycles 10-14. In both control and *crkS-RNAi* embryos, actin caps form, expand, and merge into pseudocleavage furrows. However, in *crkS-RNAi* conditions, actin cap expansion is reduced compared to controls, leading to impaired pseudocleavage furrow formation. Images acquired every 10 s and displayed at 10 frames per second. Time between frames varies between genotypes due to different sizes of region scanned. Scan time was 2.30 seconds for the control embryo and 2.93 seconds for *crkS-RNAi* embryo. Focal plane was occasionally adjusted to maintain a clear view of the furrows. This video is related to figure 6K,L.

Supplemental Movie 3. Crk loss impairs actin cap expansion and myosin remodeling during syncytial nuclear cycles. Time-lapse movies showing myosin dynamics in control and *crkS-RNAi* embryos during nuclear cycles 10-14, via MRLC(Sqh)-GFP. In control embryos, inter-cap myosin zones are remodeled during actin cap expansion, resulting in an enrichment of myosin in pseudocleavage furrows. In *crkS-RNAi* embryos, this remodeling is impaired, resulting in broader inter-cap myosin zones and reduced enrichment of myosin in pseudocleavage furrows. Images acquired every 10 s and displayed at 10 frames per second. Time between frames varies between genotypes due to different sizes of region scanned. Scan time was 2.01 seconds for the control embryo and 2.43 seconds for the *crkS-RNAi* embyo. Focal plane was occasionally adjusted to maintain a clear view of the furrows. This video is related to figure 6Q-T.

**Table 1: Description of the *Drosophila melanogaster* stocks used in this study and their sources.**

| Fly stocks | Use/Description | Source |
|---|---|---|
| *y[1], sc[*], v[1]; P{y[+t7.7] v[+t1.8]=TRiP.HMC03964}attP40* (stock #55277) | *crkS-RNAi* | Bloomington Drosophila Stock Center (Bloomington, IN, USA) |
| *y[1], sc[*], v[1]; P{y[+t7.7] v[+t1.8]=TRiP.HMJ2295}attP40/CyO* | *crkW-RNAi* | National Institute of Genetics (Mishima, Japan) |
| *w; P(mat-tub-Gal4)mat67; P(mat-tub-Gal4)mat15* (referred to as *matII; matIII*) | double-copy *mat*GAL4 | Peifer lab stocks |
| *y,w* | wild-type control | Peifer lab stocks |
| *w[*]; P{w[+mC]=His2Av-mRFP1}II.2* (stock #23651) | internal wild-type control | Bloomington Drosophila Stock Center (Bloomington, IN, USA) |
| *w[*]; P{w[+mC]=His2Av-mRFP1}III.1, P{w[+mC]=sGMCA}3.1* (stock #59023) | live imaging of F-actin dynamics (Moesin::GFP) | Bloomington Drosophila Stock Center (Bloomington, IN, USA) |
| *w[1118]; P{w[+mC]=sqh-GFP.RLC}3* (stock #57145) | live imaging of myosin dynamics (MRLC::GFP) | Bloomington Drosophila Stock Center (Bloomington, IN, USA) |
| *P{w[+mC]=ovo-FLP.R}M1A, w[*]* (stock #8727) | germline FLPase for making germline clones | Bloomington Drosophila Stock Center (Bloomington, IN, USA) |
| *y,w; crk[ΔattP]/In(4) ci[D] ci[D] pan[ciD]* | *crk* null allele | See Materials and Methods |
| *y,w; P{w[+]=mNG::3XFLAG::Crk}crk[ΔattP]* | cDNA-based rescue construct at *crk* locus | See Materials and Methods |
| *y,w; P{w[+]=FRT-mNG::3XFLAG::Crk-FRT}crk[ΔattP]* | FRT-flanked cDNA-based rescue construct at *crk* locus | See Materials and Methods |
| *y,w; P{w[+]=FRT-crk genomic-FRT}crk[ΔattP]* | FRT-flanked genomic rescue construct at *crk* locus | See Materials and Methods |
| *w; P{w[+]=UASp-mNG::3XFLAG::Crk}attP16* | mNG::3XFLAG::Crk overexpression | See Materials and Methods |
